# Transcriptomic and physiological responses of soybean plants subjected to a combination of water deficit and heat stress under field conditions

**DOI:** 10.1101/2025.05.07.652738

**Authors:** Ranjita Sinha, María Ángeles Peláez-Vico, Sameep Dhakal, Abdul Ghani, Ronald Myers, Mohit Verma, Ben Shostak, Andrew Ogden, C. Bennet Krueger, Jose R. Costa Netto, Sara I. Zandalinas, Trupti Joshi, Felix B. Fritschi, Ron Mittler

## Abstract

Water deficit, heat stress, and a combination of water deficit and heat stress are highly disruptive to crop yield worldwide. Unfortunately, the frequency and intensity of these conditions is gradually increasing due to climate change. Previous studies of water deficit and heat stress combination were primarily conducted under controlled growth conditions, revealing that the combination of water deficit and heat stress resulted in the activation of unique stress responses and acclimation pathways. However, whether similar responses to stress combination occur in the field remained largely unknown. Here we report on a two-year field study in which the transcriptomic and physiological responses of vegetative and reproductive tissues of soybean (*Glycine max*) to water deficit, heat treatment and their combination were studied. Our findings reveal that the transcriptomic responses of soybeans grown in the field are different from those grown under controlled growth chamber conditions. These differences were especially noticeable in plants subjected to the heat or water deficit treatments, and less in plants subjected to the stress combination. In addition, we report that differential transpiration between leaves and pods, that was originally discovered in plants grown under controlled growth conditions, occurs in field grown soybeans in response to heat stress, as well as heat stress combined with water deficit. We hope that the transcriptomic datasets generated by our study will contribute to future studies of crop responses to different stresses in the field, as well as highlight the need for more omics studies of plants grown under field conditions.

## INTRODUCTION

Crop plants growing in the field are subjected to a myriad of different environmental conditions that constantly change. These include light intensity, temperature, humidity, soil water content, nutrient availability, and/or the presence of different pathogens, insects, and/or herbivores. These conditions are very different from the controlled growth chamber or greenhouse conditions used in most scientific studies in which almost all environmental conditions are controlled except one or two conditions (the treatment) that are altered. As the true test of crop success in producing biomass and/or yield occurs in the field, it is important to study how crop plants respond to stress under field conditions (Mittler, 2006; Mittler and Blumwald, 2010; Nelissen et al., 2014, 2019; Cohen et al., 2021a; Rivero et al., 2022; De Meyer et al., 2023; Rhee et al., 2024; Zandalinas et al., 2024).

We previously studied the response of Arabidopsis (*Arabidopsis thaliana*), tobacco (*Nicotiana tabacum*), citrus (Carrizo citrange; *Citrus sinensis × Poncirus trifoliata*), and soybean (*Glycine max*) to a combination of water deficit and heat stress conditions (Rizhsky et al., 2002, 2004; Koussevitzky et al., 2008; Zandalinas et al., 2016, 2017; Cohen et al., 2021b). This stress combination is experienced by many crops around the world and can occur even in the absence of a true drought, at midday when temperatures rise, and soil water content is insufficient to support plant cooling by transpiration (Mittler, 2006; Mittler and Blumwald, 2010). In our controlled studies of water deficit and heat stress combination (conducted in growth chambers and/or greenhouses) we used physiological, molecular and omics approaches and discovered that the response of plants to this stress combination is unique and cannot be extrapolated from simply summing up the individual responses of plants to water deficit or heat stress (Rizhsky et al., 2002, 2004; Koussevitzky et al., 2008; Zandalinas et al., 2016, 2017; Cohen et al., 2021b; Zandalinas and Mittler, 2022). This was especially obvious in omics studies in which hundreds to thousands of different transcripts were uniquely altered in their expression only in response to the stress combination (and not in response to water deficit or heat stresses). Our findings were further supported by other studies using additional plants and crops including sorghum and barley (*e.g.,* Pradhan et al., 2022; Kumar et al., 2023; Mikołajczak et al., 2023; Ali et al., 2025).

Recently, we discovered that the molecular and physiological responses of reproductive tissues of soybean (flowers and pods) are different from those of vegetative tissues (leaves) and that during a combination of water deficit and heat stress leaf stomata of soybean plants were closed and transpiration suppressed, while at the same time, stomata of flowers and pods (of the same plants) remained open and transpiration from these organs continued. This process, termed ‘differential transpiration’ (between vegetative and reproductive tissues), enabled the transpirational cooling of reproductive tissues during the stress combination and protected these tissues from heat related damages (Sinha et al., 2022, 2023a, 2023b, 2025). Differential transpiration was subsequently reported to also occur in tomato confirming our findings with soybean (Jensen et al., 2024).

Although a combination of water deficit and heat stress was extensively studied using growth chamber and/or greenhouse conditions in different plants (Rizhsky et al., 2002, 2004; Koussevitzky et al., 2008; Zandalinas et al., 2016, 2017; Cohen et al., 2021a, 2021b), and multiple studies have shown that it has an overall negative impact on crop yield (*e.g.,* Cohen et al., 2021a), it is largely unknown whether this stress combination has the same impact on the transcriptomics and physiological responses of plants growing in the field. In addition, it is unknown whether differential transpiration occurs in soybean plants subjected to a combination of water deficit and heat stress under field conditions. To address these questions, we conducted a field experiment in the summers of 2022 and 2024 in which soybean plants growing in the field were subjected to no additional stress (control), water deficit, heat treatment, and a combination of water deficit and heat treatment. The two main hypotheses we wanted to test using these studies were: *i*) that plants grown in the field will have a unique response to different stress/stress combination conditions that might not always be replicated using growth chamber conditions, and *ii*) that differential transpiration occurs under field conditions. In addition, we aimed to generate transcriptomic datasets from field grown plants that could serve as a comparative resource for the scientific community to test whether the findings identified in the laboratory indeed happen under field conditions.

## RESULTS AND DISCUSSION

### Subjecting soybean plants to water deficit, heat treatment and their combination under field conditions

To induce conditions of water deficit, heat stress, and water deficit combined with heat stress in soybean plants (*cv Magellan*) grown in the field, we used a combination of mobile rainout shelters, controlled irrigation, and heat treatment provided by infra-red heaters (Figure 1a-1c). Control plants were maintained well-watered and grown alongside water deficit-, heat treated-, and water deficit plus heat-treated plants using a split plot design (Figure S1a). Soil water potential and soil temperature were measured for control and water deficit plots using soil sensors (Figure 1d), and air humidity, vapor pressure deficit (VPD), and temperature were constantly monitored for the entire research area (Figure 1a) using a fully automated weather station (Figures 1e, S1b). Field experiments were conducted at the University of Missouri Bradford Research Center (38°53′N, 92°12′W) in 2022 and 2024. In 2022, plants were planted on May 13^th^, water deficit stress was initiated on June 10^th^, and heat treatment (for heat treatment and water deficit plus heat treatment) was initiated on July 25^th^. Plants were then sampled on July 30^th^ and conditions at the time of sampling are shown in Figures 1, S1b, and Table S1. In 2024, plants were planted on May 23^rd^, water deficit stress was initiated on August 1^st^ (later than in 2022 due to rain), and heat treatment (for heat treatment and water deficit plus heat treatment) was initiated on August 19^th^. Plants were then sampled on August 23^rd^ and conditions at the time of sampling are shown in Figures 1, S1b, and Table S1.

**Figure 1.**
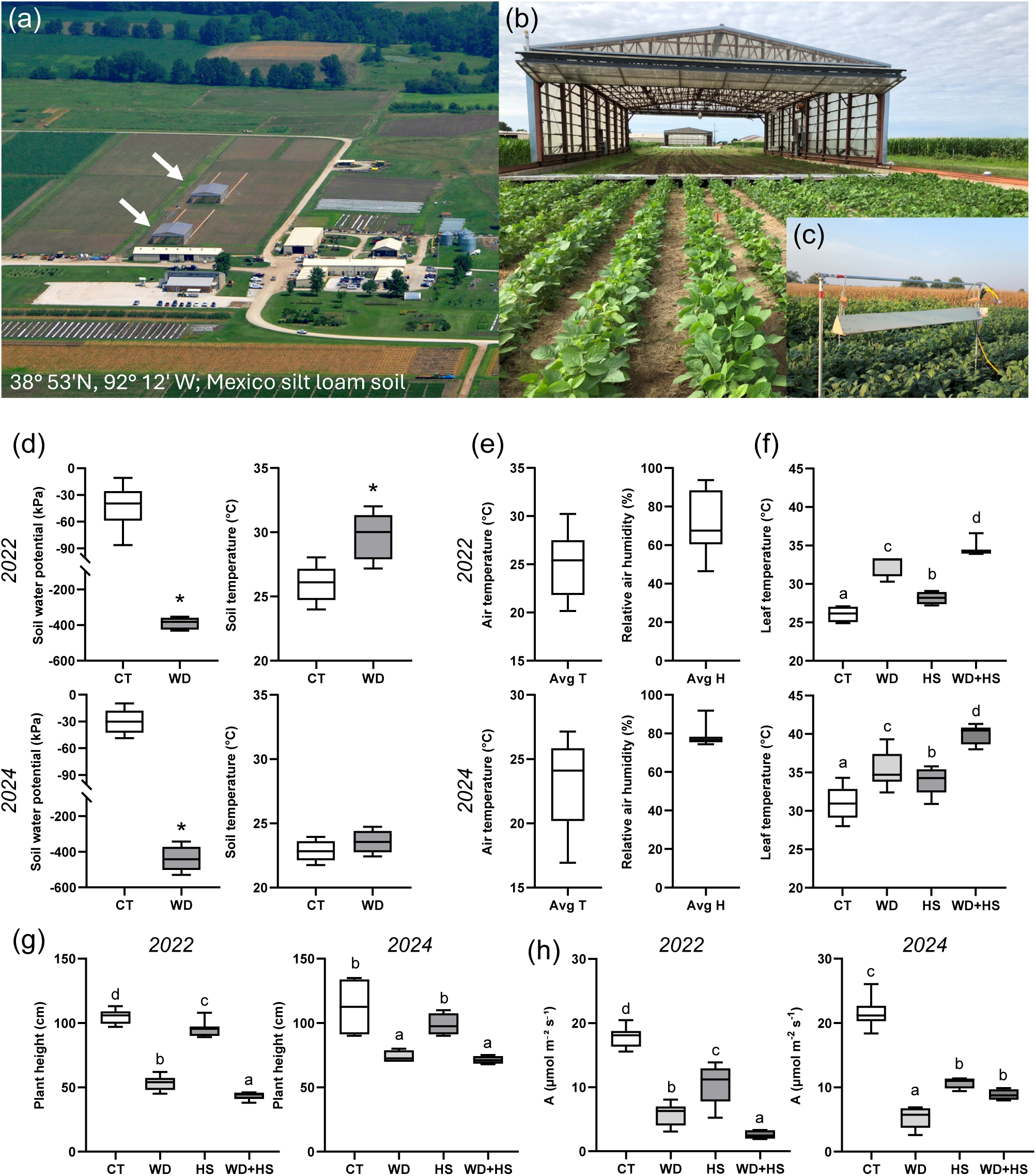
Applying a combination of water deficit and heat stress under field conditions. Field experiments were conducted during the growth season in 2022 and 2024 at the Bradford research center, Columbia, Missouri, US. (a) An aerial image showing the two rainout shelters used for the experiments (white arrows). (b) An image showing rows of soybean growing for the experiment and the rail system used to mobilize the rainout shelters. (c) An image of an infrared heater hanging above the plants. (d) Average soil water potential and temperature for the month of July at the two years. (e) Same as in d but for air temperature and humidity. (f) Leaf temperature at the time of sampling for 22 and 24. (g) Plant height measured a week after the last sampling on 22 and 24. (h) Photosynthetic activity at the time of sampling for 22 and 24. Box plots represent an average of at least 12-15 measurements from three split plot replicates. Student’s t-test was used for testing significance differences between control and water-deficit in panel (d). Asterisk (*) denotes significant difference at P < 0.05. One-way-ANOVA with Fisher’s least square difference (Fisher’s LSD) was used as test of significance for panels (f) to (h). Different letters denote significant difference between means at P < 0.05. Additional environmental parameters at the field site before, during, and following sampling are shown in Figure S1b. Abbreviations: 22, 2022 sample; 24, 2024 sample; A, CO_2_ assimilation; Avg, average; CT, control; HS, heat treatment; WD, water-deficit; WD + HS, a combination of WD and HS.

Leaf temperature measurements taken from the different plants using an infra-red camera in 2022 and 2024 revealed that, in contrast to chamber experiments conducted with soybean plants of the same genotype (Sinha et al., 2022), leaf temperature of water deficit-treated plants was higher than that of heat-treated plants in the field (Figure 1f). In contrast, and in agreement with the previous chamber experiments (Sinha et al., 2022), leaf temperature of soybean plants subjected to a combination of water deficit and heat treatment in 2022 and 2024 was higher than that of plants subjected to water deficit or heat treatment (Figure 1f). The differences in leaf temperature between the individual water deficit and heat stress treatments in the field (Figure 1f) and the growth chamber experiments (Sinha et al., 2022), were likely a result of differences in soil water availability. As opposed to plants grown in pots in the growth chamber, root growth of heat-treated and irrigated plants in the field was not restricted which could have allowed for greater water uptake and transpirational cooling. Moreover, water deficit-treated plants in the field may have been subjected to higher temperatures than growth chamber plants, while at the same time lacked sufficient water to cool their leaves. In support of this possibility, the leaf water potential of heat treated plants in the field (Figure S2; measured in 2022) was higher than that of heat stress plants treated in the growth chamber (Sinha et al., 2022), while the leaf water potential of water deficit treated plants grown in the field (Figure S2; measured in 2022) was much lower compared to growth chamber grown plants (Sinha et al., 2022). In addition, field ambient temperatures and canopy level temperatures easily reached 36-42 ^ο^C during midday (Figure S1b for 2022 and 2024, and S3 measured in 2022, respectively), while growth chamber temperature was stable at 38 ^ο^C for heat-treated plants, and 28 ^ο^C for water deficit treated plants (Sinha et al., 2022).

The impact of the different stress treatments on soybean plants grown in the field was further determined by measuring changes in plant height (Figure 1g) and leaf photosynthesis (Figure 1h). These measurements revealed that water deficit, and water deficit combined with heat treatment, had a significant impact on plant growth and photosynthetic activity. Compared to leaf photosynthetic measurements of water deficit-treated plants (of the same soybean genotype) under growth chamber conditions (Sinha et al., 2022), the impact of water deficit on photosynthesis was more robust under field conditions (Figure 1h). Like the effects of water deficit on leaf temperature measurements (Figure 1f), this result could also be an outcome of limited water availability under field conditions coupled with higher temperatures (compared to the controlled conditions used in the chamber experiments; Figures S1b, S2, S3; Sinha et al., 2022). Similar to water deficit and water deficit combined with heat treatment, heat treatment applied alone also resulted in suppressed photosynthesis (Figure 1h), but only slightly suppressed growth (Figure 1g). These findings agree with those observed in the chamber studies (Sinha et al., 2022). Taken together, the measurements presented in Figures 1, and S1b, S2, S3 demonstrate that the setup we used in the field induced conditions of water deficit, heat stress and a combination of water deficit and heat stress in soybean plants grown in the field, and that the combination of water deficit and heat treatment had an overall negative impact on plant growth (Figure 1g).

### Differential transpiration under field conditions

Differential transpiration in soybean was discovered and studied in plants grown in growth chambers (Sinha et al., 2022, 2023b). To determine whether differential transpiration also occurs in field grown soybeans, we measured transpiration and stomatal conductance of soybean leaves (vegetative tissue) and pods (reproductive tissue) from plants grown in the field, and subjected to control, water deficit, heat treatment and a combination of water deficit and heat treatment (Figures 1, 2, S1). In response to water deficit, and in agreement with the chamber experiments, transpiration and stomatal conductance were suppressed in both leaves and pods (compared to control; Figure 2). In contrast, in response to heat treatment and compared to control, transpiration and stomatal conductance of leaves were significantly suppressed, while transpiration and stomatal conductance of pods were unchanged, revealing that differential transpiration can occur in field grown soybeans under heat conditions applied in the absence of any induced water deficit (Figure 2). This result was different from the growth chamber experiments in which transpiration and stomatal conductance of leaves was maintained high under heat stress conditions (Sinha et al., 2023b; potentially a result of differences in VPD and ambient temperature between the growth chambers and the field environments). In response to a combination of water deficit and heat treatment, and in agreement with all previous studies conducted in chambers (Sinha et al., 2022, 2023b), compared to control, transpiration and stomatal conductance of leaves were significantly suppressed, while transpiration and stomatal conductance of pods were unchanged (*i.e.,* differential transpiration; Figure 2). The findings presented in Figure 2 reveal therefore that differential transpiration between leaves and pods occurs in field grown soybeans under a combination of water deficit and heat stress (similar to the experiments conducted in chambers; Sinha et al., 2023b). Interestingly, in the field (but not in the chambers), compared to control, differential transpiration also occurred in soybean plants subjected to heat treatment alone (in the absence of induced water deficit; Figure 2). This finding could reflect the higher temperatures, VPD, and/or the potential for lower water availability that occur in the field, compared to the controlled conditions in the chambers. Taken together, our findings reveal that differential transpiration between leaves and pods can occur in field grown plants under conditions of heat stress, and/or heat stress combined with water deficit. As heat stress applied during the reproductive stages of soybean causes significant reduction in yield (*i.e.,* Siebers et al., 2015; Djanaguiraman et al., 2019; Thomey et al., 2019; Soba et al., 2022), cooling of pods by differential transpiration (Figure 2) could play a key role in protecting yield from heat and water deficit plus heat stresses in the field.

**Figure 2.**
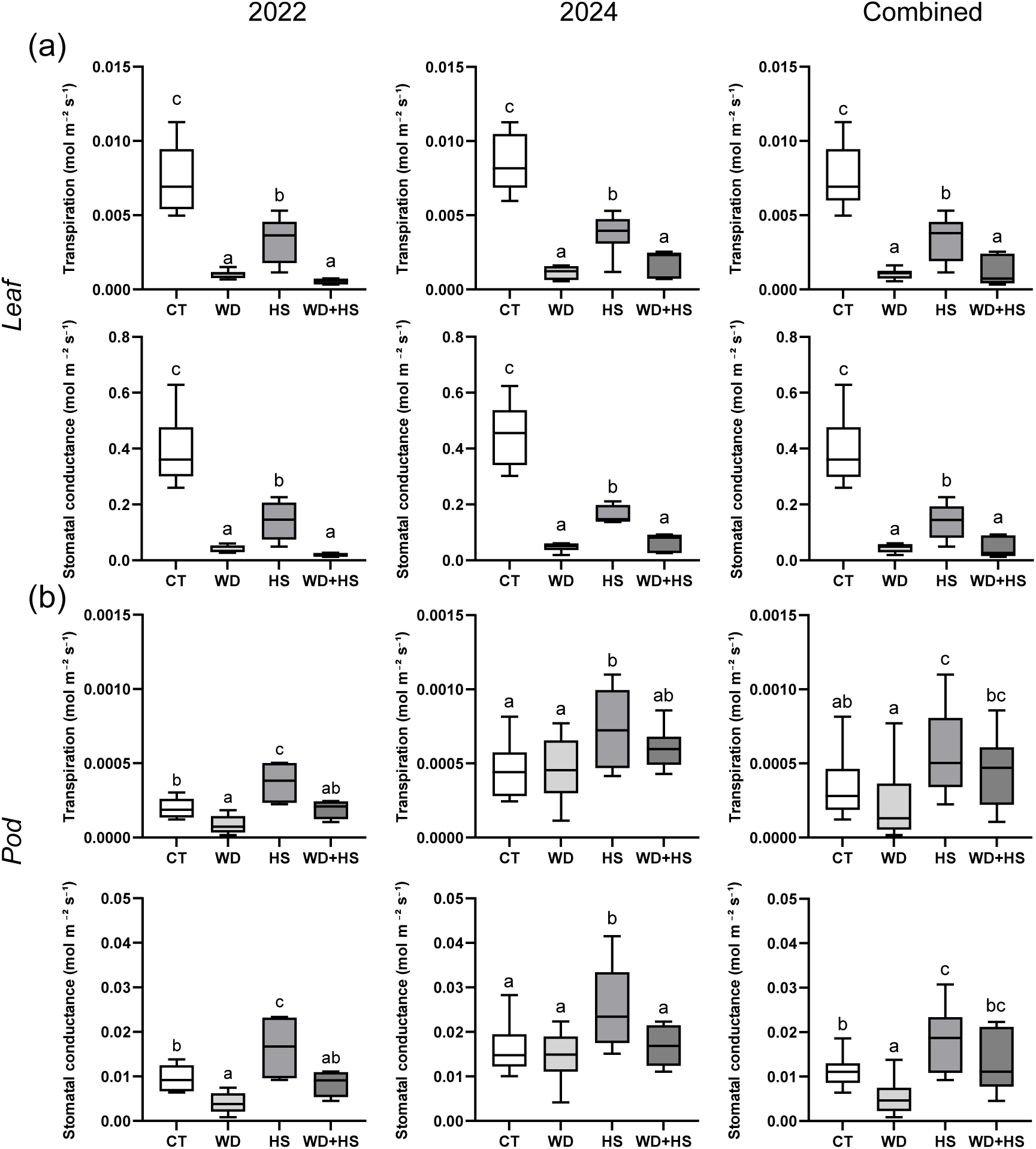
Differential transpiration between leaves in pods in soybean plants grown in the field. Transpiration and stomatal conductance of soybean leaves (a) and pods (b) were measured using LICOR Portable Photosynthesis System during 2022 and 2024 (24) (Figure S1b). Plants were grown in the absence of stress (CT), under water deficit (WD), heat treatment (HS), or a combination of WD+HS under field conditions as described in Figures 1 and S1. Box plots represent an average of at least 12-15 measurements from three split plot replicates. One-way-ANOVA with Fisher’s least square difference (Fisher’s LSD) was used to test for significance. Different letters denote significant difference at P < 0.05. Abbreviations: 22, 2022 sample; 24, 2024 sample; CT, control; HS, heat treatment; WD, water-deficit; WD + HS, a combination of WD and HS.

### Transcriptomic analysis of soybean plants subjected to water deficit, heat stress and their combination under field conditions

The transcriptomic responses of vegetative and reproductive tissues of soybean plants to water deficit, heat stress, and a combination of water deficit and heat stress were previously studied using growth chamber conditions (Sinha et al., 2022, 2023a, 2023b). These studies revealed that the transcriptomic response of soybeans to a combination of water deficit and heat stress is different than that to water deficit or heat stress, and that marked differences exist between the response of reproductive and vegetative tissues to the stress combination (Sinha et al., 2022, 2023a, 2023b). Based on the analysis described above, certain pathways were identified as pathways of interest to be used to enhance differential transpiration. These include abscisic acid (ABA) degradation and stomatal number regulation in reproductive tissues, and heat stress responses in both vegetative and reproductive tissues (Sinha et al., 2022, 2023a, 2023b, 2025). However, to what extent the transcriptomic responses to stress combination that occur in the growth chamber are similar to those that occur in the field is unknown. To begin addressing this question, we conducted RNA-Seq analysis of soybean leaves and pods sampled during 2022 and 2024 (designated 22 and 24; Figure S1b) from field grown soybeans, without any added stress (control), or subjected in the field to water deficit, heat treatment, and their combination (Figures 1-3, S1; Tables S2-S7).

**Figure 3.**
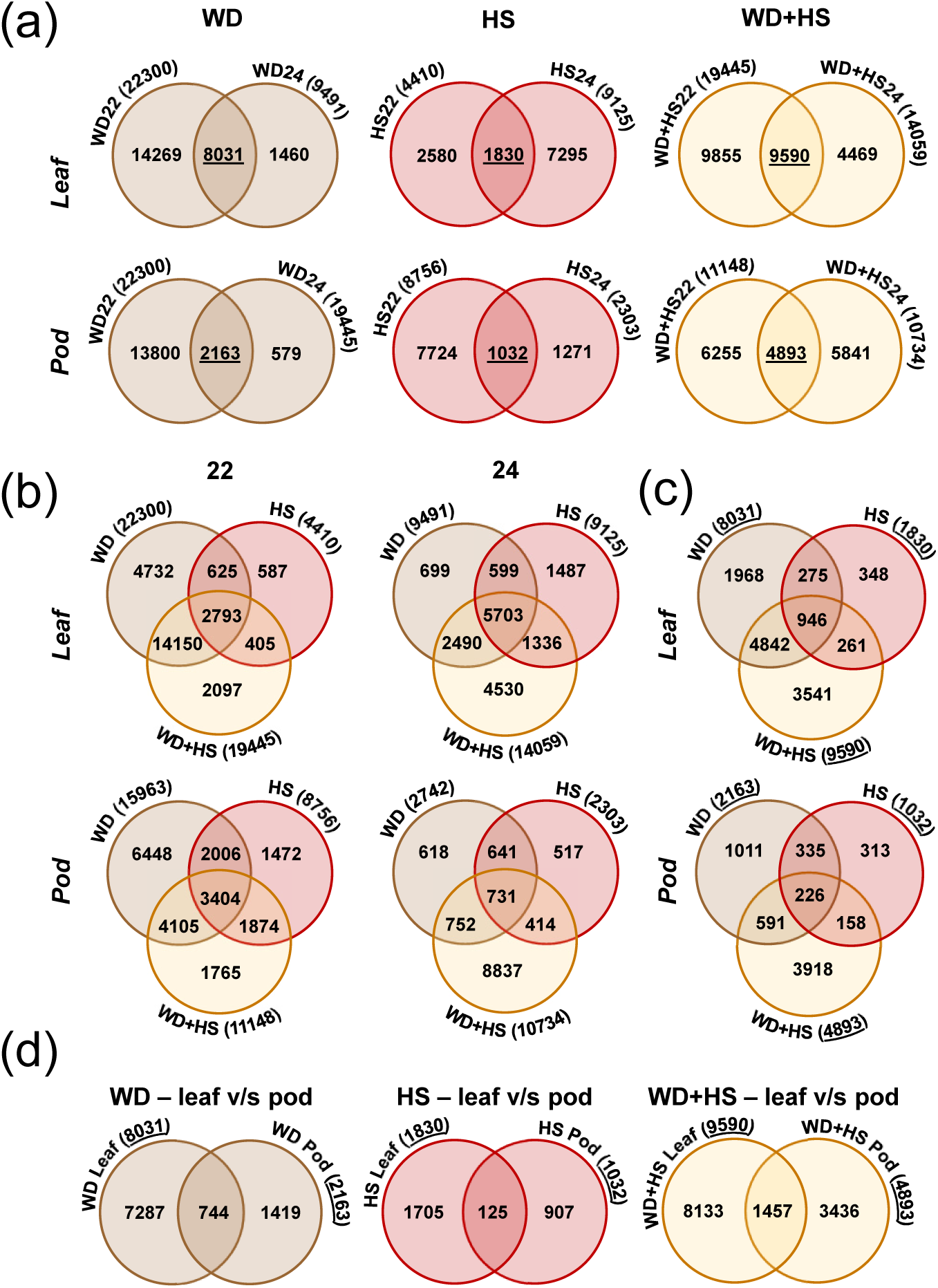
Transcriptomics responses of soybean plants grown in the field and subjected to water deficit, heat stress and water deficit plus heat stress combination. Leaf and pod transcripts with a differential expression pattern (significantly enhanced or suppressed) between control (CT) and water deficit (WD), heat treatment (HS), or a combination of WD+HS were identified. (a) Venn diagrams showing the overlap between all WD-, HS-, or WD+HS-altered transcripts sampled in 22, and 24, respectively. Overlap between leaf transcripts is shown on top and overlap between pod transcripts is shown on bottom. (b) Venn diagrams showing the overlap between WD-, HS-, and WD+HS-altered transcripts in each sampling time (22 and 24, respectively). Overlap between leaf transcripts is shown on top and overlap between pod transcripts is shown on bottom. (c) Venn diagrams showing the overlap between all 22, and 24 conserved WD-, HS-, and WD+HS-altered transcripts (from a; underlined). (d) Venn diagrams showing the overlap between leaf and pods conserved (from a; underlined) WD-, HS-, and WD+HS-altered transcripts. Significant changes in transcript expression compared to control were defined as adjusted P < 0.05 (negative binomial Wald test followed by Benjamini–Hochberg correction). Abbreviations: 22, 2022 sample; 24, 2024 sample; CT, control; HS, heat stress; WD, water-deficit; WD + HS, a combination of WD and HS.

As expected from field experiments that involve dynamic conditions and multiple and mostly uncontrolled variables, a comparison between the leaf and pod transcriptomic responses of soybean to each of the different stress conditions (water deficit, heat treatment, or a combination of water deficit and heat treatment), between the different sampling times/years (22 and 24; Figure S1b), revealed marked differences between each sampling time/year (Figure 3a). These differences were also reflected in a KEGG pathway enrichment analysis of the different leaves and pods transcriptomics data from the different sampling times (Figures S4). For example, the KEGG pathways enriched in leaves from heat treated plants (as well as pods from water deficit treated plants) were highly variable between 22 and 24 (Figure S4).

As shown in Figure 3b, and despite the differences in the response of soybean to each individual stress treatment at each sampling time/year (Figures 3a, S4), the leaf and pod transcriptomic responses of field grown soybean plants subject to the water deficit and heat treatment stress combination contained many unique transcripts that were not altered in their expression in response to water deficit or heat (confirmed for all measurements, conducted in two different years). The identification of thousands of transcripts uniquely expressed during the stress combination (Figures 3b, S4) was similar to that observed in the chamber experiments (Sinha et al., 2022, 2023a, 2023b).

As each sampling time/year displayed such a large variability in transcript expression (Figure 3a), we repeated the analysis shown in Figure 3b, but this time used the conserved transcripts identified for each treatment in Figure 3a (underlined). As shown in Figure 3c, a comparison between the conserved leaf and pod transcriptomic responses to water deficit, heat treatment, or their combination (from Figure 3a) yielded similar results to the analysis shown in Figure 3b, demonstrating that the conserved transcriptomic responses of leaf and pods to the stress combination are also unique (Figure 3c).

A gene ontology (GO) analysis of the transcripts common to all stresses in leaves (Figure 3c; 946 transcripts) identified many transcripts involved in photosynthesis, heat stress response, and reactive oxygen species (ROS) metabolism (Figure S5). In contrast, a GO analysis of the transcripts unique to the stress combination in leaves (Figure 3c; 3541 transcripts) identified transcripts involved in carboxylic acid metabolism, thiolester hydrolase activity, and plastid organization (Figure S6). Interestingly, a GO analysis of the transcripts common to all stresses in pods (Figure 3c; 226 transcripts), identified transcripts involved in oxidoreductase activity, hypoxia, and wounding (Figure S7). In contrast, a GO analysis of the transcripts unique to the stress combination in pods (Figure 3c; 3918 transcripts) identified transcripts involved in translation, histone modification, and mRNA binding (Figure S8). Taken together, the GO analyses of the common and unique transcripts to pods and leaves subjected to the stress combination in the field (Figures 3c, S5-S8) revealed that leaves expressed more transcripts involved in heat stress and ROS responses, while pods expressed more transcripts involved in sugar metabolism, oxidoreductases and hypoxia.

We further used the conserved transcriptomic responses from Figure 3a to conduct a comparison between the transcriptomic responses of leaves and pods in the field to the different stresses (Figure 3d). At least in growth chamber experiments, the transcriptomic responses of leaves and pods were found to be very different from each other (Sinha et al., 2023a, 2023b). As shown in Figure 3d, and in agreement with the chamber experiments (Sinha et al., 2023a, 2023b), the conserved responses of pods and leaves to each of the stress conditions were different from each other.

Taken together, the findings presented in Figure 3 reveal that like growth chamber grown plants (Sinha et al., 2023a, 2023b), the leaf and pod responses of field grown plants to the stress combination are unique, but different from each other. However, how similar are the responses of growth chamber and field stressed soybean plants is an open question that is addressed below.

### Comparing the transcriptomic responses of soybean plants to water deficit, heat treatment and their combination between field and growth chamber conditions

To identify similarities and differences between the growth chamber and field transcriptomic responses of soybean plants to water deficit, heat treatment and their combination, we reprocessed the RNA-Seq data previously obtained for soybean plants subjected to these treatments in growth chambers (Sinha et al., 2022, 2023a, 2023b) together with our current field RNA-Seq data (Figure 3). We then compared the leaf and pod transcripts significantly altered in their expression in response to water deficit, heat treatment, and their combination in the field (conserved among the different seasons/samples; underlined; Figure 3c) to all transcripts significantly altered in their expression in response to the same types of stress conditions applied in the growth chamber (Sinha et al., 2022, 2023a, 2023b), keeping in mind the differences and similarities between the two environments, described above (Figures 1, 2, S1).

To gauge the degree of overlap between the transcriptomics responses of soybean plants (of the same genotype) grown in the field or the growth chamber and subjected to water deficit, heat treatment and their combination, we determined how many of the transcripts identified as significantly altered in their expression in the field were also identified as significantly altered in the growth chamber, in leaves and pods, in response to the different stress conditions. As shown in Figure 4a, a relatively high degree of overlap between the growth chamber and the field transcriptomic responses was found in leaves and pods subjected to the stress combination (35% and 48% of the conserved field transcripts were also detected in leaves and pods from chamber experiments, respectively). In contrast, the responses of leaves and pods to all other stress conditions were less similar (about 24% and 6% of the conserved water deficit leaf and pods field transcripts were also detected in tissues from the chambers, and about 22% and 23% of the conserved heat stress leaf and pod field transcripts were also detected in chamber-grown plants; Figure 4a).

**Figure 4.**
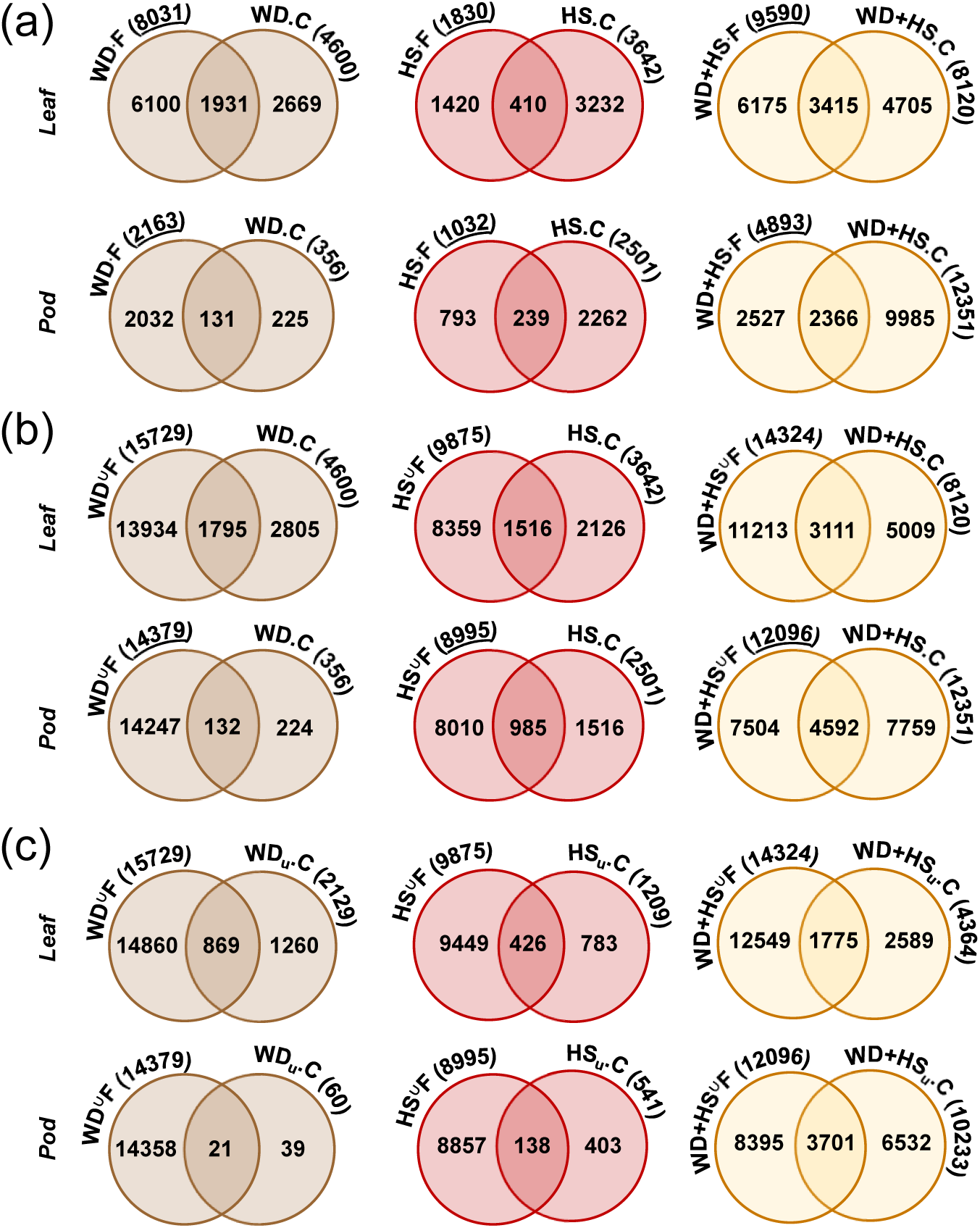
Similarities and differences between the transcriptomic responses of soybean grown in the field or the growth chamber and subjected to water deficit, heat stress and water deficit plus heat stress combination. Transcriptomics data for field and growth chamber plants was processed together, and leaf and pod transcripts with a differential expression pattern (significantly enhanced or suppressed) between control (CT) and water deficit (WD), heat treatment (HS), or a combination of WD+HS were identified in field or growth chamber grown plants. (a) Venn diagrams showing the overlap between WD-, HS-, and WD+HS-altered transcripts from plants grown in the field (conserved transcripts from Figure 3c) and all WD-, HS-, and WD+HS-altered transcripts from growth chambers. Overlap between leaf transcripts is shown on top and overlap between pod transcripts is shown on bottom. (b) Same as in a, but for unique field WD-, HS-, and WD+HS-altered transcripts from the field (unique to each batch year/time pooled together; represented as **^∪^**) and all WD-, HS-, and WD+HS-altered transcripts from growth chambers. (c) Same as in b, but comparing unique field WD-, HS-, and WD+HS-altered transcripts from the field (unique to each batch year/time pooled together; represented as **^∪^**) and differentially expressed transcripts unique to each stress treatments in the chambers (represented as u*). Significant changes in transcript expression compared to control were defined as adjusted P < 0.05 (negative binomial Wald test followed by Benjamini–Hochberg correction). Abbreviations: C, chamber; CT, control; F, field; HS, heat treatment; **^∪^**, union of unique transcripts; u, unique; WD, water-deficit; WD + HS, a combination of WD and HS.

As expected, less overlap was observed when the unique responses to each stress condition in the field were compared to all transcripts significantly altered in their expression in response to the same types of stress conditions applied in the growth chamber (Figure 4b). An exception was the degree of similarity between the unique responses of pods to the stress combination (about 37% of unique field transcripts were detected by the chamber experiments; Figure 4b). Low overlap was also observed when the unique responses to each stress condition in the field were compared to the unique responses to each stress in the growth chamber (Figure 4c). Overall, compared to the leaf and pod transcriptomic responses to water deficit or heat treatment, that were relatively different between the field and the chamber environments, the transcriptomic responses of leaf and pods to the combination of water deficit and heat treatment (both conserved and unique) was less diverged between the field and the chamber (Figure 4a, 4b). This finding suggests that unlike water deficit or heat treatment that could have multiple and redundant response pathways (and hence induce different responses in the growth chamber and the field), the combination of water deficit and heat treatment may impose a much more constrained type of stress on plants that induces a less diverged and/or redundant set of pathways (and hence induces more similar responses in the field and the chamber). More studies are required to address this intriguing possibility.

To provide an initial (albeit limited) understanding to the differences between the responses of soybean in the field and the growth chamber, we conducted a GO analysis of the transcripts that are unique to the field in the response of soybeans to the different stress conditions. We initially conducted this analysis on the conserved transcripts unique to the field (Figures 4a, S9-S14) and then extended it to the unique transcripts from the field experiments in all seasons (combined) compared to unique transcripts from growth chamber experiment (Figures 4c, S15-S20). These analyses revealed that transcripts involved in photosynthesis were enriched in the response of leaves to all stress conditions in the field (Figures S9-S11). Transcripts encoding iron-sulfur proteins and oxidoreductases were further enriched in leaves of plants subjected to water deficit in the field, while transcripts encoding sugar phosphate and organic-acid related functions, as well as iron-sulfur proteins, were further enriched in leaves from plants subjected to water deficit plus heat treatment in the field (Figures S9-S11). Pods from plants subjected to water deficit in the field contained a high proportion of transcripts encoding cytoskeleton, cell wall, and DNA binding proteins (Figure S12). In contrast, the transcriptome of pods subjected to heat stress in the field was enriched in transcripts encoding oxidoreductases and cell wall biosynthesis (Figure S13). Transcripts enriched in RNA binding and protein biosynthesis functions were further found to accompany the transcriptomic response of pods to the combination of water deficit and heat stress in the field (Figure S14). An examination of the GO classification of transcripts unique to the field environment in both leaves and pods (from Figure 4c) revealed that RNA binding, ribosome biogenesis, and translation were highly represented in all these samples (Figures S15-S20).

Taken together, the findings presented in Figure 4 and Figures S9-S20 reveal that the transcriptomic responses of soybeans to different stress conditions in the field differed considerably from those of growth chamber grown plants. Given the varied conditions and degree of stress experienced by the plants in the different environments, these differences were not surprising. Nevertheless, they were more pronounced in plants subjected to the heat treatment or water deficit applied alone and less pronounced in plants subjected to the water deficit and heat treatment combination. In addition, our analysis revealed that certain groups of transcripts (*e.g.,* those involved in photosynthesis or ribosome assembly/translation) are enriched in field grown plants.

Further studies are needed to dissect the transcriptomic responses obtained in the field and the datasets reported here should facilitate such future studies that are critical for our ability to generate climate resilient crops. Below, we will focus on two of the pathways important for heat stress tolerance and differential transpiration, *i.e.,* heat shock transcription factors (HSFs) and ABA biosynthesis/degradation, respectively.

### Expression of transcripts involved in ABA biosynthesis/degradation, heat stress regulation, and circadian/rhythmic processes in pods and leaves of field and growth chamber grown plants

The HSF network of plants regulates many plant responses to stress and could be used as an excellent indicator of plant stress levels/responses (Jacob et al., 2017; Bakery et al., 2024). We therefore used the expression level of soybean HSFs to compare between the field and growth chamber experiments. As shown in Figure 5a, except for some HSFs (*e.g.,* GLYMA_14G096800 and GLYMA_01G185800) in leaves, changes in the expression of HSFs mostly did not correlate between the field and the growth chamber environments. Moreover, only a few HSFs had a similar expression pattern in all seasons/samples from the field (and these did not generally match the growth chamber experiments; Figure 5a). Taken together, our findings reveal that changes in the expression level of some HSFs occurred only under field conditions, and that the expression pattern of many HSFs did not correlate between the different seasons/samples. These findings could reflect the unique conditions that occur in the field in each season/time and highlight the need to study plants in the field (as opposed to controlled growth chamber/greenhouse). In addition, they could be a result of time-of-day sampling, reflecting a need for multiple sampling times over hourly and daily cycles.

**Figure 5.**
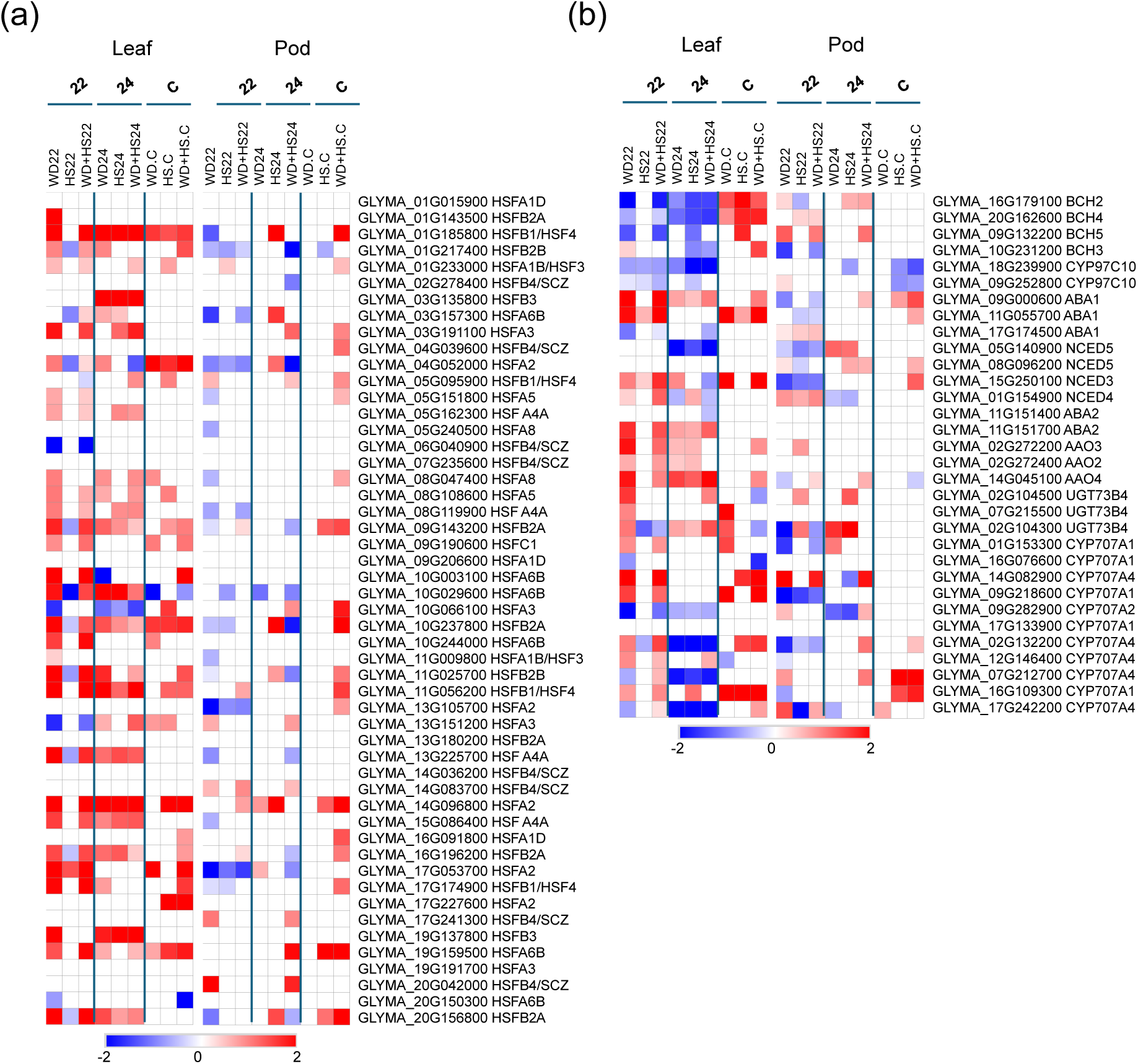
Expression of transcripts involved in abscisic acid (ABA) metabolism and heat stress regulation in field and chamber grown plants subjected to a combination of water deficit and heat stress. Heat maps showing the differential expression of transcripts encoding heat shock transcription factors (HSFs) (a), or enzymes involved in ABA biosynthesis, degradation, and conjugation (b), in field and growth chamber grown plants subjected to water deficit, heat treatment, and their combination are shown. Significant changes in transcript expression compared to control were defined as adjusted P < 0.05 (negative binomial Wald test followed by Benjamini– Hochberg correction). Abbreviations: ABA, abscisic acid; C, chamber; HS, heat treatment; WD, water-deficit; WD + HS, a combination of WD and HS.

Using controlled growth chamber conditions, we previously reported that differential transpiration between pods and leaves, or flowers and leaves, is associated with enhanced ABA degradation, as well as expression of ABA degrading enzymes (Sinha et al., 2022, 2023a, 2023b). We therefore determined whether a similar pattern of transcript expression is also found in leaves and pods from plants grown in the field. As shown in Figure 5b, the elevated expression of *CYP707A1* and *CYP707A4* (*ABA 8’-hydroxylase*; GLYMA_16G109300 and GLYMA_07G212700) that occurred in pods of growth chamber grown plants subjected to the combination of water deficit and heat treatment (Sinha et al., 2023a) did not consistently occur in pods from different seasons/samples. Instead, the expression of a different *CYP707A4* (GLYMA_14G082900) was consistent in pods from plants subjected to the water deficit and heat treatment combination in the field (Figure 5b). This finding suggests that ABA degradation in pods, a process that is important for differential transpiration (Sinha et al., 2022, 2023a, 2023b, 2025), could be regulated differently in field and growth chamber grown plants. Another notable difference between the chamber and field experiments was the expression of *BCH2* (*beta carotene hydroxylase 2*) that was elevated in leaves from growth chamber grown plants but was mostly suppressed in leaves from the field grown plants. This finding could suggest that ABA/beta carotene biosynthesis in leaves of field grown plants is regulated differently than in chamber grown plants (Sinha et al., 2022, 2023a, 2023b).

While comparing the field and chamber transcriptomics responses, we observed marked differences in these two groups in the expression pattern of transcripts involved in rhythmic and circadian processes. These were especially notable in leaves (Figure 6) but also occurred in pods (Figure S21). As circadian and rhythmic processes are affected by changes in environmental conditions (Legris et al., 2017; Xiong et al., 2022; Tiwari et al., 2023), it is likely that the fluctuating light, temperature, humidity, and/or other conditions in the field (compared to the more stable conditions in the chamber), impacted the expression of these transcripts. A protein-protein interaction-based network analysis of the different genes shown in Figures 6a and 6b revealed that flowering time associated genes (*i.e.,* GLYMA_18G027500; *EARLY FLOWERING 4b*; center in the map in panel c) could be playing a key role in regulating circadian and rhythmic processes in the field (Figure 6c). As soybean *cv Magellan* is indeterminant when it comes to flowering, and synchronized growth and flowering in soybean are dependent on environmental conditions (*i.e.,* Tian et al., 2010; Kim et al., 2022), the high representation of circadian and rhythmic processes in field grown plants and their link to flowering is most likely a result of fluctuating light and temperature conditions in the field (*e.g.,* fluctuating light intensity/quality due to cloud cover and canopy movements, as well as fluctuating temperature and humidity due to changes in precipitation, wind speed, and wind direction), compared to the steady conditions of the growth chamber. Further studies are needed to determine how plants respond to such fluctuating conditions in the presence or absence of stress and how these interactions impact flowering and yield in the field. Taken together, the findings presented in Figures 3-6 support the hypothesis that plants grown in the field will have a highly complex and specialized response to different environmental stress conditions, that cannot always be reliably replicated or predicted using growth chamber conditions.

**Figure 6.**
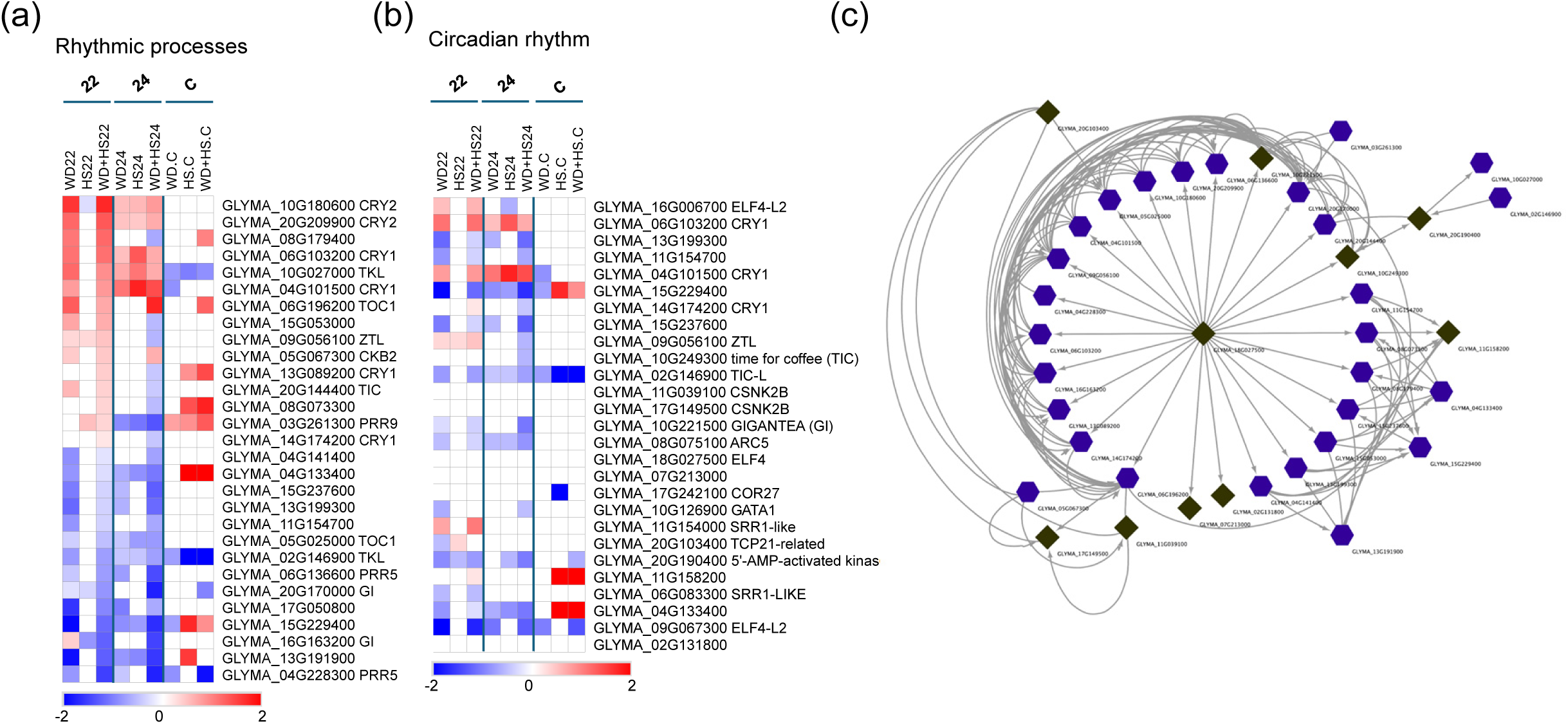
Expression of transcripts involved in rhythmic processes, flowering time and circadian rhythm in field and chamber grown plants subjected to a combination of water deficit and heat stress. Heat maps showing the differential expression of transcripts encoding proteins involved in flowering time and rhythmic processes (a), and circadian rhythm (b), in field and growth chamber grown plants subjected to water deficit, heat treatment, and their combination are shown together with a protein-protein interaction-based network of the proteins shown in a and b (c). Nodes represent genes (or proteins), while edges indicate known interactions. Specific node shapes and colors were used to distinguish key functional groups: Hexagonal-shaped nodes represent core genes belonging to rhythmic process; Diamond-shaped nodes represent core genes belonging to Circadian rhythm; Directionality, arrow point from the regulator to the regulated gene/protein. Significant changes in transcript expression compared to control were defined as adjusted P < 0.05 (negative binomial Wald test followed by Benjamini–Hochberg correction). Abbreviations: C, chamber; HS, heat treatment; WD, water-deficit; WD + HS, a combination of WD and HS.

## Summary

Our work reveals marked differences between the transcriptomic responses of soybean plants grown in the field or the growth chamber to different stress conditions and their combination (Figures 3-6). These differences should be taken into consideration when developing genetically modified crops with enhanced resilience to climate change, as the true test of resilience and yield improvement under stress conditions occurs in the field and not in the growth chamber or greenhouse (Mittler and Blumwald, 2010; Nelissen et al., 2014; Rhee et al., 2024). We hope that the datasets generated by our study will contribute to future studies of crop responses to stress in the field, as well as highlight the need for more molecular and omics studies of crops grown under field conditions. Our study further reveals that differential transpiration occurs in the field (Figure 2), but that compared to the growth chamber environment, it may be controlled by different ABA metabolizing genes (Figure 5b). Conducting more studies on field grown plants is essential if we want to generate climate resilient crops that will allow us to meet the food demands of our growing population in the face of the stresses imposed on agriculture by global warming, climate change, and increased environmental pollution (Long et al., 2015; Rillig et al., 2019; Alizadeh et al., 2020; Lesk et al., 2022; Lee et al., 2023).

## EXPERIMENTAL PROCEDURES

### Field experiments and stress application

Field experiments were conducted at the University of Missouri Bradford Research Center (Missouri Agricultural Experiment Station, Columbia, Missouri, USA, 38°53′N, 92°12′W; Figure 1a). Soybean (*Glycine max*) plants (*cv Magellan*; Maturity Group IV, relative maturity 4.3. Indeterminate growth habit) were grown on Mexico silt loam (fine, smectitic, mesic Vertic Epiaqualf) soil during the growing seasons of 2022, and 2024 (May to September). Plants were grown in a 30 m x 15 m area that can be covered by a mobile rainout shelter (Figure 1b). The experiment was laid out as a split-plot design with water treatment as main plot and heat treated as sub-plot (Figure S1a). Soybean were planted on May, 13^th^ in 2022 and May, 23^rd^ in 2024 in rows that were 6.1 m long and 0.76 m apart from each other at a density of 34.5 seed m^-2^ as described by (Arifuzzaman et al., 2023; Costa Netto et al., 2025).

Water deficit treatments were imposed by a combination of rainout shelter deployment and irrigation (Sen et al., 2023). Control plants were supplemented with irrigation to maintain well-watered conditions throughout the experiments. Heat treatment was applied as described by (Kimball, 2005; Cavanagh et al., 2022) using infrared heaters (Kalglo®-Infrared Heater HS-2420 I 65” I 240V I 2000W, Fogelsville, PA, USA) installed 30 cm above the canopy (Figure 1c). A combination of water deficit and heat treatment was applied by installing the heaters under the mobile rainout shelters. No differences were found between the effects of space or infrared heaters on stomatal aperture (measured as explained in Sinha et al., 2022) and tissue temperature of pods and leaves in preliminary experiments conducted in 2022 (Figure S22).

Experimental conditions consisted of control conditions (Plants were irrigated to maintain soil water potential at 0.15 m soil depth at > – 40 kPa), water deficit stress (A combination of rainout shelters and irrigation leading to an average soil water potential at 0.15 m soil depth of < –100 kPa under ambient conditions), heat treatment (A combination of irrigation to soil water potential of > – 40 kPa and heat provided by the infrared heaters), and water deficit and heat treatment combination (Plants exposed to a soil water potential of < –100 kPa and supplemented with heat provided by the infrared heaters). Ambient conditions were continuously monitored using a weather station (http://agebb.missouri.edu/weather/realtime/columbiaBREC.asp), soil water potential was measured using Teros 21 soil sensors (TEROS 21 Tech Specs, METER group, Pullman, WA, USA), canopy and leaf temperature were measured with an infrared camera (FLIR C2; FLIR Systems AB, Wilsonville, OR, USA), and leaf water potential was measured using a Scholander pressure chamber (600 Pressure Chamber Instrument, PMS Instrument, Albany, OR, USA) between 11.30 AM –12.00 PM.

Soybean plants were grown in the field until the R1 developmental stage (Fehr et al., 1981) and then subjected to well-watered (control) and water deficit stress (water deficit) for 3 weeks. Heat treatment was applied to well-watered (heat treatment) and water deficit treated (water deficit plus heat treatment) plants 4-5 days before the end of the 3-week water deficit stress period (Table S1). All sampling (control, water deficit, heat treated, and water deficit plus heat treated) was conducted 3 weeks post water deficit treatment initiation, aside from plant height that was measured a week after sampling for RNA-Seq analysis (Table S1).

### Gas exchange parameters

Stomatal conductance, transpiration, and photosynthesis of fully developed leaves and developing pods (2-3 cm size) from the 3^rd^ node (from top) of the different plants were measured using a Li-Cor Portable Photosynthesis System (LI-6800; Li-Cor, Lincoln, NE, USA) between 10:00 and 12:00 h, as described by (Sinha et al., 2022). Measuring window for photosynthesis and other physiological parameters was determined by studying the physiological responses of crops at 10:00, 12:00, and 14:00 during the 2022 season (Figure S23).

### RNA-Seq analysis

Soybean leaves and pods were collected from plants grown in the field under control, water deficit stress, heat treatment, and water deficit and heat treatment combination between 10:30 and 11:30 h and immediately frozen in liquid nitrogen (Figure S1b). Fully developed leaves and developing pods (2-3 cm size) were collected from the 3^rd^ node (from top) of each plant. For 2022 batch (22), samples were collected during the 4^th^ week of July (Figure S1b) and for the 2024 batch (24), samples were collected during the 4^th^ week of August 2024 (Figure S1b). Out of 4 split plot replicates, 3 random split plots were used as 3 biological repeats (Figure S1a). For each technical repeat, leaves and pods at the same developmental stage (Sinha et al., 2023a) were pooled from 3-6 plants, and each biological repeat consisted of 3 technical repeats. RNA was isolated using RNAeasy Plant Mini Kit (Qiagen). RNA libraries for sequencing were prepared using standard Illumina protocols and RNA sequencing was performed by Novogene Co. Ltd (Sacramento, CA, USA; https://en.novogene.com/) using NovaSeq 6000 PE150.

For RNA-Seq analysis, raw RNA-Seq data from the two batches: Batch 22, and Batch 24, were processed together with raw RNA-Seq data from previous chamber experiments (Sinha et al., 2022, 2023a, 2023b) using a Python-based analysis pipeline developed to process raw sequencing data. This analysis generated a list of genes along with FPKM values for each batch. Then, differential expression between control, water deficit, heat, and water deficit plus heat was determined for the two field batches, and for the chamber experiments, separately and differentially expressed genes for each batch were determined. The workflow involved checksum verification to ensure data integrity, quality assessment of individual sequencing read using FastQC v0.12.1 (https://www.bioinformatics.babraham.ac.uk/projects/fastqc/) and comprehensive quality report compilation of sequencing files with MultiQC. Further, potential adapter contamination from raw data was eliminated using TrimGalore v0.6.10 (https://www.bioinformatics.babraham.ac.uk/projects/trim_galore/), followed by re-evaluation using FastQC v.0.12.1 and MultiQC to confirm data quality and the effective removal of adapter sequences. Prior to further analysis, the gene model and reference genome were downloaded from Phytozome (https://phytozome-next.jgi.doe.gov/). Reference genome Gmax_880_v6.0, and gene model Gmax_880_Wm82.a6.v1, comprising a total of 48,387 genes was used for further analysis. Processed reads were subsequently aligned to the reference genome using Hisat2 (Kim et al., 2019), achieving alignment rates above 97% for Batch 22, and above 88% for Batch 2024 across individual pairs of FASTQ files. The resulting alignment files were saved in BAM format and further processed using Samtools v1.18 (Danecek et al., 2021) for sorting. The sorted BAM files were used for transcript assembly and quantification to generate FPKM and differential expression using Cufflinks v2.2.1 (Trapnell et al., 2012), The genes.diff file obtained from Cuffdiff output (Trapnell et al., 2013) was used to identify significant genes, defined as those with a q-value ≤ 0.05. Venn diagrams were constructed with Venny 2.0 (BioinfoGP, CNB-CSIC), Heat maps were generated in Morpheus (https://software.broadinstitute.org/morpheus) and overrepresented gene ontology (GO) terms (P < 0.05) and KEGG pathway enrichment (P < 0.05) were conducted using g:profiler (Raudvere et al., 2019) and ShinyGO 0.82 (Ge et al., 2020; https://bioinformatics.sdstate.edu/go/), respectively. Dot-plot for enriched KEGG pathways were created in R using ggplot2. RNA-Seq data is available at Gene Expression Omnibus (GEO) under accession GSE290497.

### Network Construction

A protein-protein interaction (PPI) network was constructed for the proteins encoded by the genes shown in Figure 6a and 6b using STRING v12.0 (https://string-db.org/; Szklarczyk et al., 2023). The gene set of interest was uploaded to STRING, and interactions with a minimum confidence score of 0.4 (medium confidence) were retrieved. Only experimentally validated interactions and those derived from curated databases were considered. The resulting interactions data were exported in TSV format for further analysis. The visualization layout was optimized using the yFiles Radial Layout algorithm for better interpretation of interaction patterns, particularly through the yFiles Layouts plugin. The exported files were further imported into Cytoscape v3.10.3 (Shannon et al., 2003) for visualization and network analysis.

## DECLARATION OF AI USE

AI-assisted technologies were not used in creating this article.

## AUTHOR CONTRIBUTIONS

RS, MAP-V, AG, RJM, SD, MV, BS, AO, BK, JRCN performed the experiments and analyzed the data. RM, FBF, TJ, RS, MAP-V, and SIZ designed experiments, analyzed the data, and/or wrote the manuscript.

## ACKNOWLEDGMENTS

We thank the support and services provided by the University of Missouri College of Agriculture Food and Natural Resources and the Bradford Research Center.

## DATA AVAILABILITY STATEMENT

RNA-Seq data were deposited in the Gene Expression Omnibus (GEO) database, under the following accession numbers: GSE290497.

## SUPPORTING INFORMATION

Additional Supporting Information may be found in the online version of this article.

## SUPPLEMENTAL FIGURES

**Figure S1.** Experimental design and environmental conditions at the time of sampling.

**Figure S2.** Leaf water potential of field and chamber grown plants subjected to water deficit, heat stress, and a combination of water deficit and heat stress.

**Figure S3.** Air temperature and humidity near canopy.

**Figure S4.** KEGG pathway enrichment of differentially expressed leaf (a) and pods (b) transcripts with altered expression in response to water deficit, heat treatment, and a combination of water deficit and heat treatment (compared to control) in 22, and 24.

**Figure S5.** GO enrichment of differentially expressed leaf transcripts overlapping between WD-, HS-, and WD+HS.

**Figure S6.** GO enrichment of differentially expressed leaf transcripts unique to WD+HS.

**Figure S7.** GO enrichment of differentially expressed pod transcripts overlapping between WD-, HS-, and WD+HS

**Figure S8.** GO enrichment of differentially expressed pod transcripts unique to WD+HS.

**Figure S9.** Gene ontology (GO) enriched categories for leaf transcripts uniquely altered in response to WD-stress in field compared to WD-stress in growth chamber.

**Figure S10.** Gene ontology (GO) enriched categories for leaf transcripts uniquely altered in response to HS-stress in field compared to HS-stress in growth chamber.

**Figure S11.** Gene ontology (GO) enriched categories for leaf transcripts uniquely altered in response to WD+HS-stress in field compared to WD+HS-stress in growth chamber.

**Figure S12.** Gene ontology (GO) enriched categories for pod transcripts uniquely altered in response to WD-stress in field compared to WD-stress in growth chamber.

**Figure S13.** Gene ontology (GO) enriched categories for pod transcripts uniquely altered in response to HS-stress in field compared to HS-stress in growth chamber.

**Figure S14.** Gene ontology (GO) enriched categories for pod transcripts uniquely altered in response to WD+HS-stress in field compared to WD+HS-stress in growth chamber.

**Figure S15.** Gene ontology (GO) enrichment of WD induced leaf transcripts unique to field compared to growth chamber.

**Figure S16.** Gene ontology (GO) enrichment of HS induced leaf transcripts unique to field compared to growth chamber.

**Figure S17.** Gene ontology (GO) enrichment of WD+HS induced leaf transcripts unique to field compared to growth chamber.

**Figure S18.** Gene ontology (GO) enrichment of WD induced pod transcripts unique to field compared to growth chamber.

**Figure S19.** Gene ontology (GO) enrichment of HS induced pod transcripts unique to field compared to growth chamber.

**Figure S20.** Gene ontology (GO) enrichment of WD+HS induced pod transcripts unique to field compared to growth chamber.

**Figure S21.** Differential expression of soybean pod transcripts enriched in circadian rhythm and rhythmic processes GO categories.

**Figure S22.** Effects of regular or infrared heaters on stomatal aperture and tissue temperature of pods and leaves.

**Figure S23.** Measurement of gas exchange parameters at different time-of-day in the field.

## SUPPLEMENTAL TABLES

**Table S1.** Experimental details and environmental parameters

**Table S2.** Leaf transcripts significantly altered in response to WD, HS and WD+HS in field experiment 2022

**Table S3.** Pod transcripts significantly altered in response to WD, HS and WD+HS in field experiment 2022

**Table S4.** Leaf transcripts significantly altered in response to WD, HS and WD+HS in field experiment 2024

**Table S5.** Pod transcripts significantly altered in response to WD, HS and WD+HS in field experiment 2024

**Table S6.** Leaf transcripts significantly altered in response to WD, HS and WD+HS in growth chamber

**Table S7.** Pod transcripts significantly altered in response to WD, HS and WD+HS in growth chamber

## SUPPORTING INFORMATION

**Figure S1.**
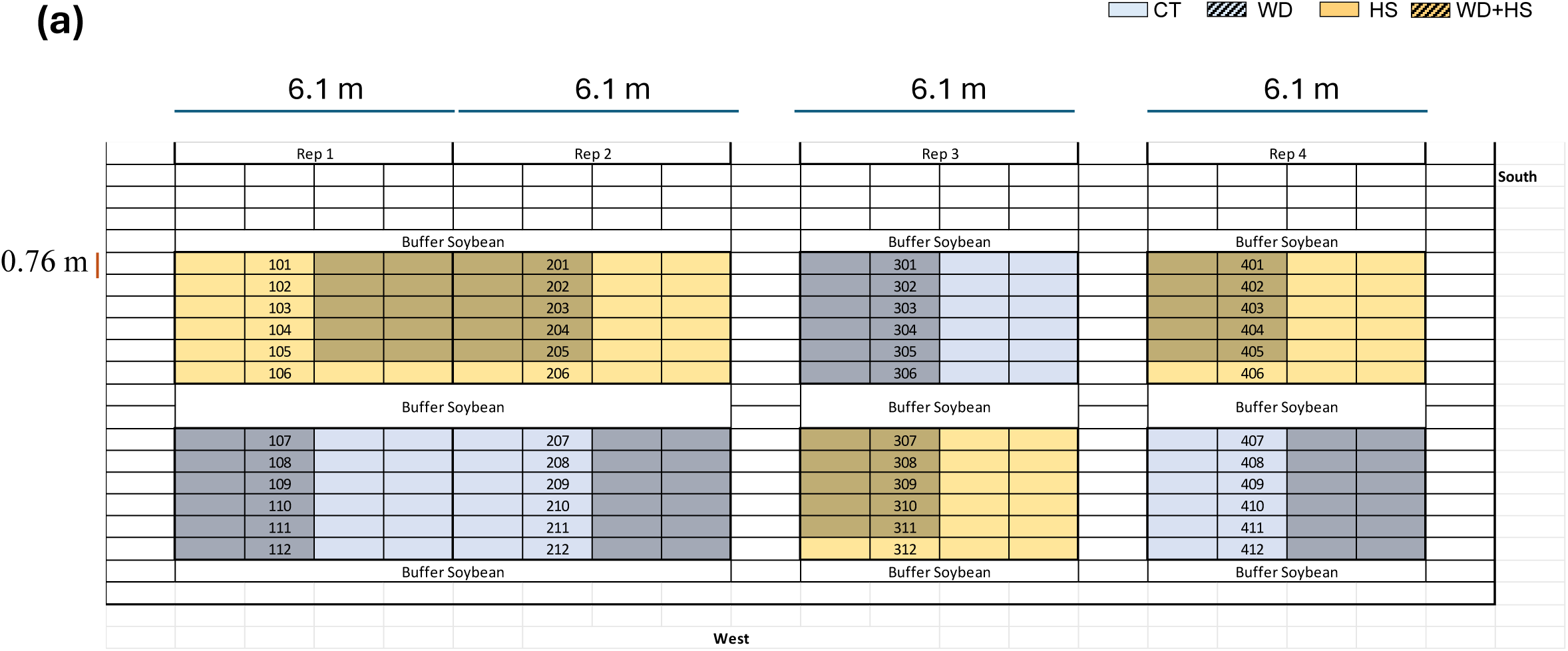
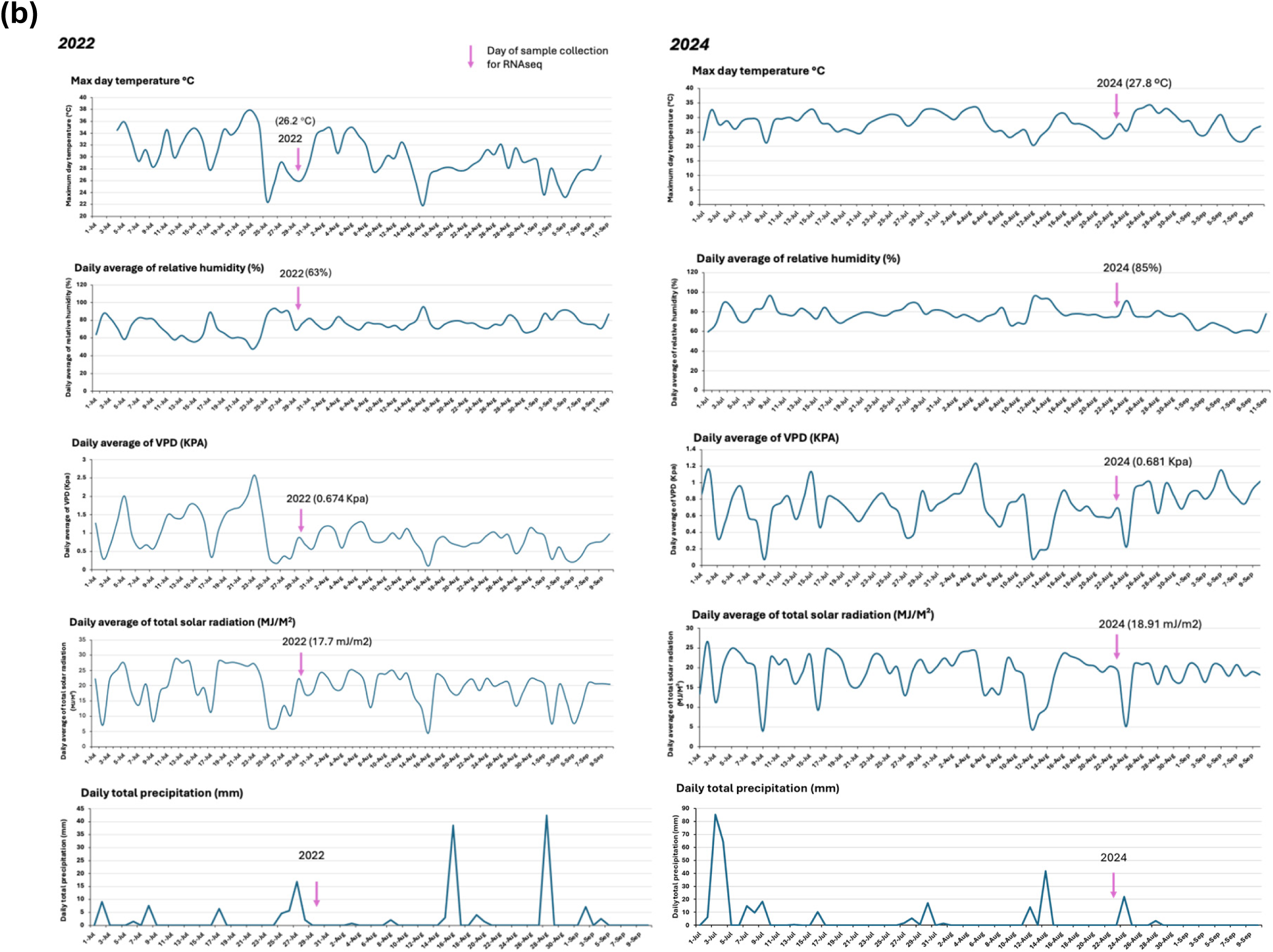
Experimental design and environmental conditions at the time of sampling. (a) The split plot experimental design used in 2022 and 2024 to subject soybean plants to water deficit, heat treatment, and a combination of water deficit and heat treatment. (b) Environmental parameters at the field prior, during, and following sampling. Numbers at the top and left side represent distance of each split plot in meter and distance between rows in meter, respectively. Abbreviations: CT, control; HS, heat stress; WD, water-deficit; WD + HS, a combination of WD and HS.

**Figure S2.**
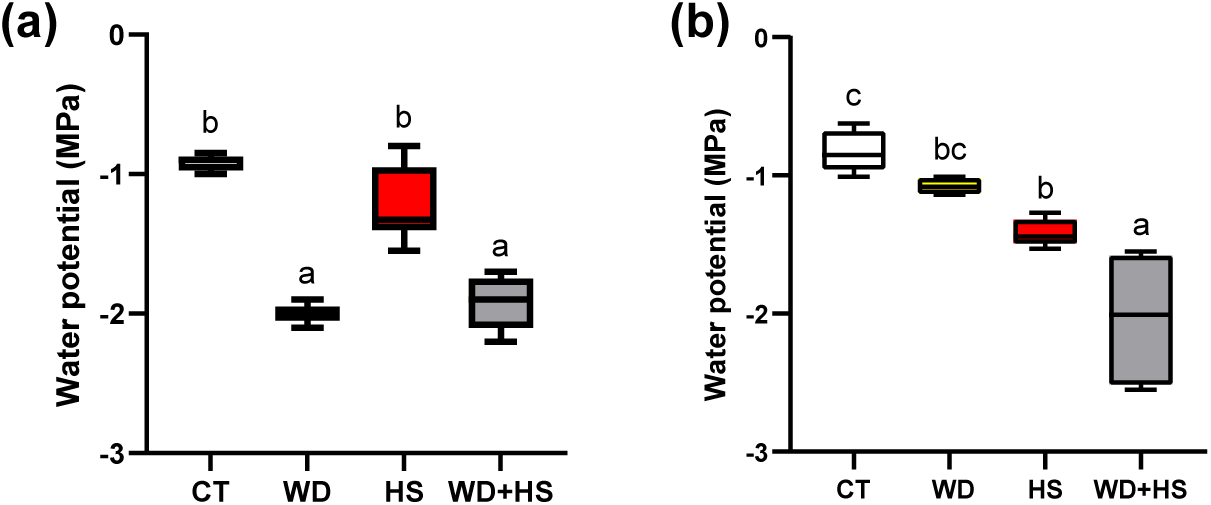
Leaf water potential of field and chamber grown plants subjected to water deficit, heat stress, and a combination of water deficit and heat stress. Leaf water potential was measured for soybean plants growing in field (a) or growth chambers (b) under conditions of water deficit, heat stress, and a combination of water deficit and heat stress. One-way-ANOVA with Fisher’s least square difference (Fisher’s LSD) was used as test of significance. Different letters denote significant difference between means at P < 0.05. Abbreviations: CT, control; HS, heat stress; WD, water-deficit; WD + HS, a combination of WD and HS.

**Figure S3.**
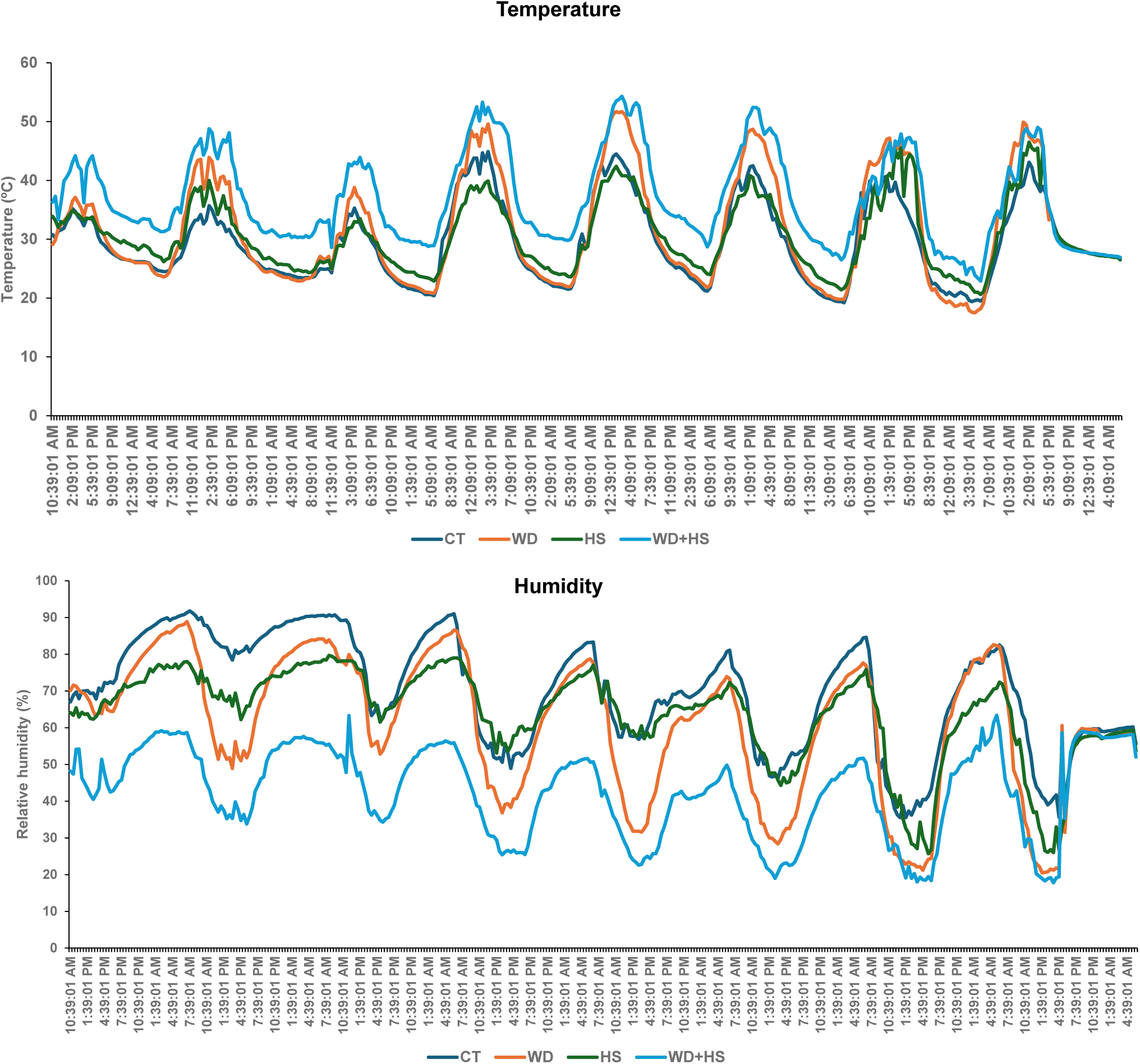
Air temperature and humidity near canopy. Air temperature and humidity were recorded near the canopy of soybean plants growing under water deficit, heat stress, and a combination of water deficit and heat stress using thermo/RH data loggers. Line graph represents temperature and humidity data of one week from July 7^th^ to July 15^th^, 2022. Abbreviations: CT, control; HS, heat stress; WD, water-deficit; WD + HS, a combination of WD and HS.

**Figure S4.**
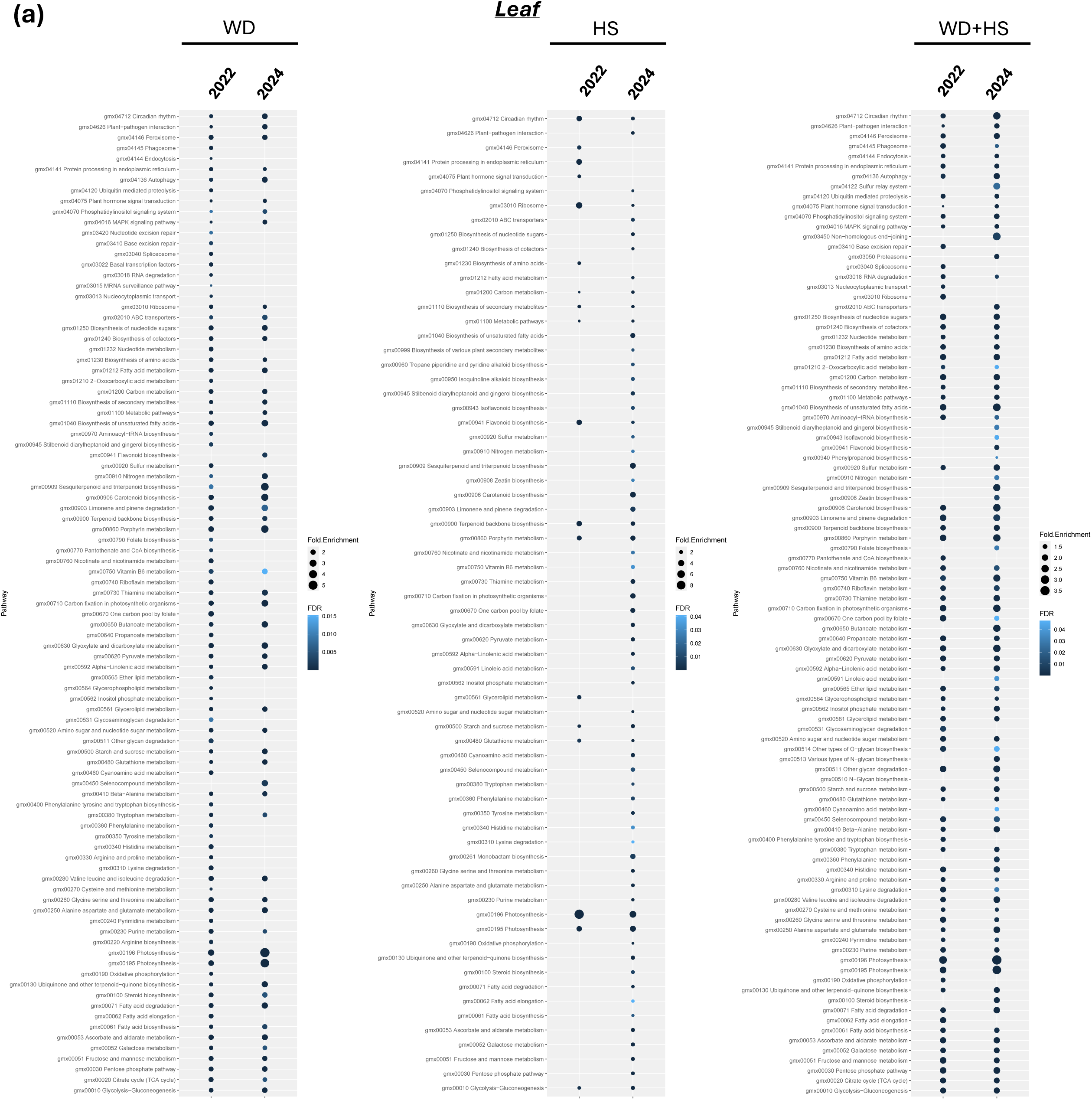
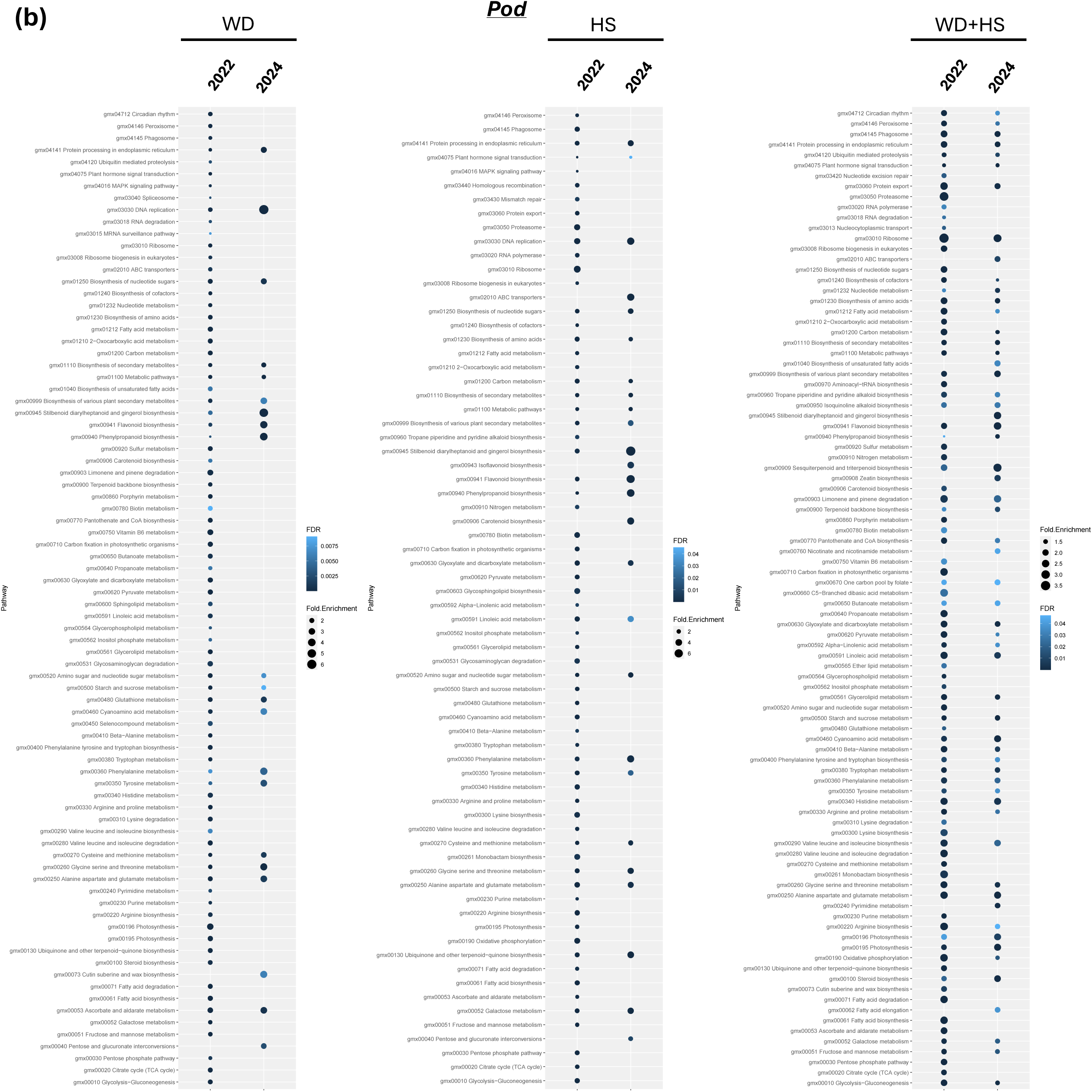
KEGG pathway enrichment of differentially expressed leaf. (a) and pods (b) transcripts with altered expression in response to water deficit, heat treatment, and a combination of water deficit and heat treatment (compared to control) in 22, and 24. Abbreviations: 22, 2022 sample; 24, 2024 sample; HS, heat treatment; WD, water-deficit; WD + HS, a combination of WD and HS.

**Figure S5.**
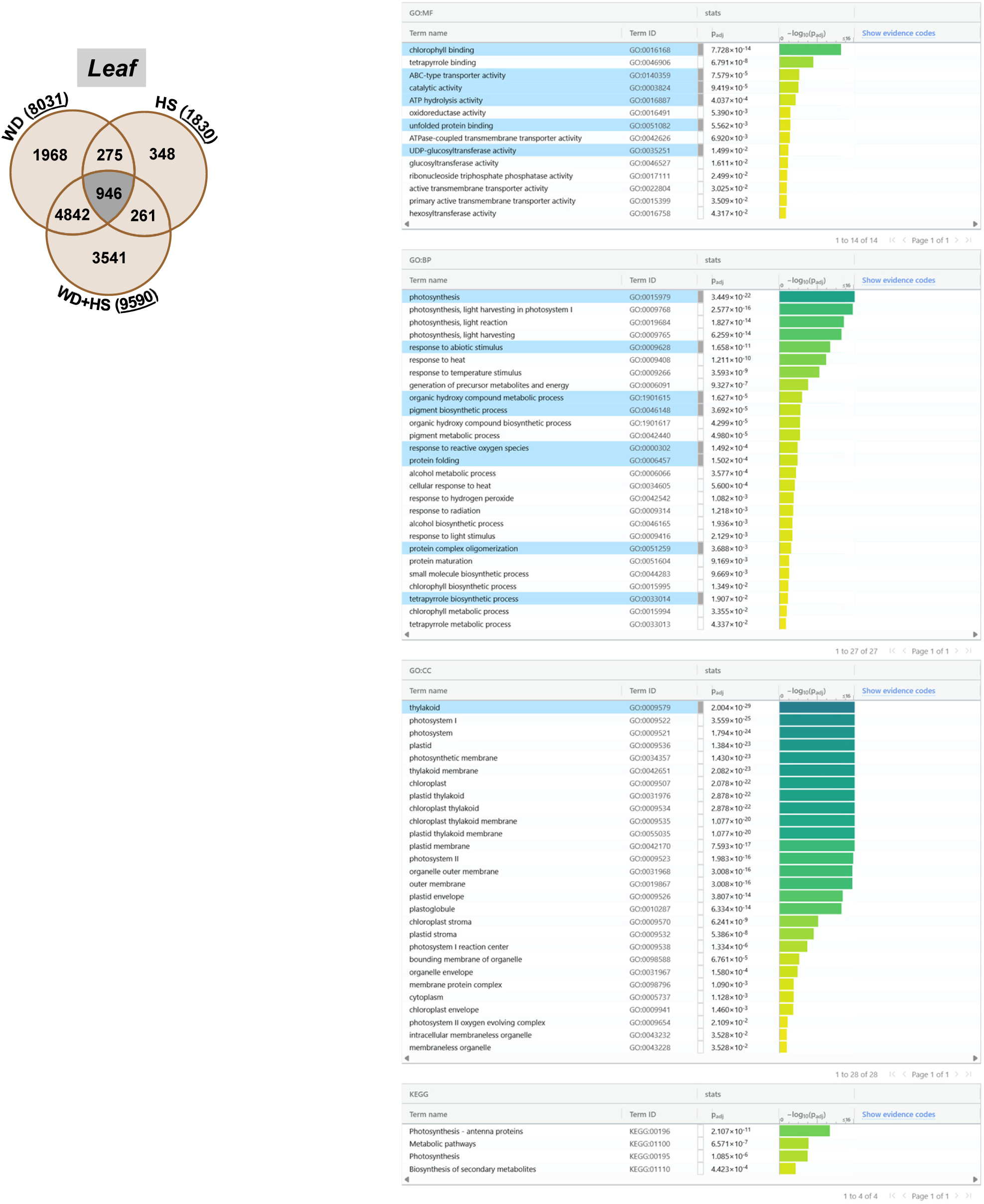
GO enrichment of differentially expressed leaf transcripts overlapping between WD-, HS-, and WD+HS. Gene ontology (GO) enriched categories for leaf transcripts overlapping between WD-, HS-, and WD+HS-altered transcripts (946 transcripts, conserved transcripts from Figure 3c). Abbreviations: BP, biological process; CC, cellular component; HS, heat stress; MF, molecular function; WD, water deficit; WD + HS, a combination of WD and HS.

**Figure S6.**
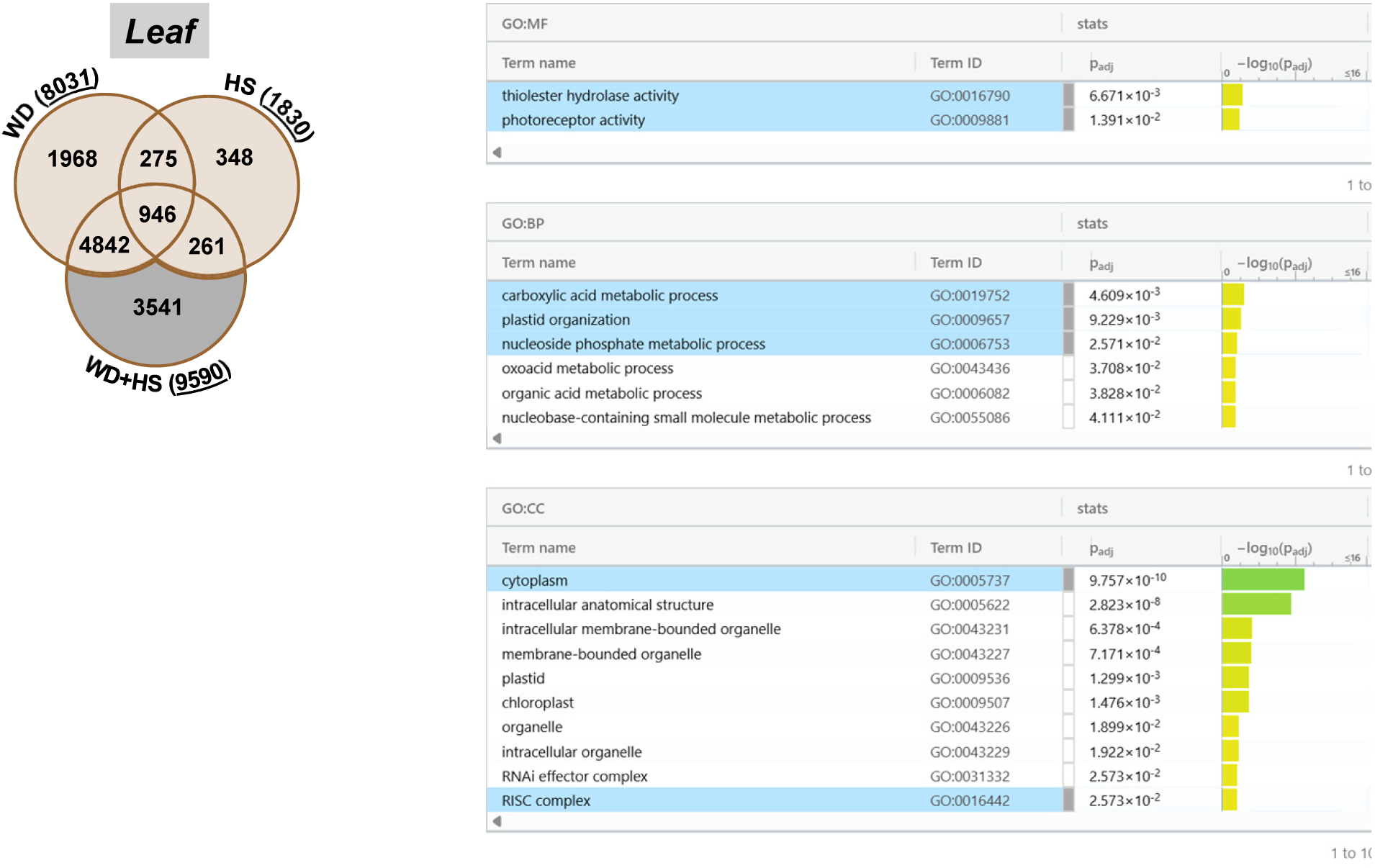
GO enrichment of differentially expressed leaf transcripts unique to WD+HS. Gene ontology (GO) enriched categories for leaf transcripts uniquely altered under WD+HS stress (3541 transcripts, conserved transcripts from Figure 3c). Abbreviations: BP, biological process; CC, cellular component; HS, heat stress; MF, molecular function; WD, water deficit; WD + HS, a combination of WD and HS.

**Figure S7.**
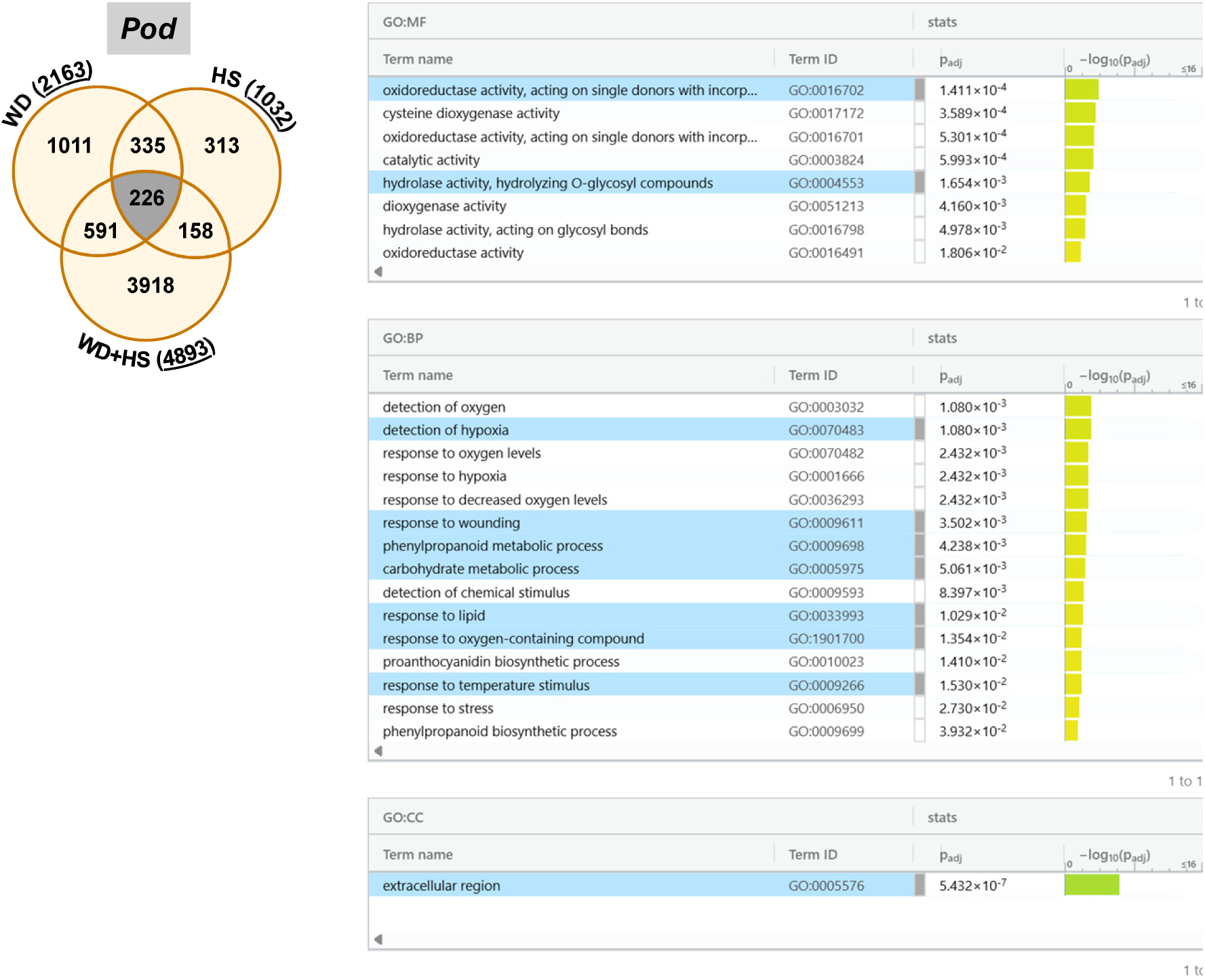
GO enrichment of differentially expressed pod transcripts overlapping between WD-, HS-, and WD+HS. Gene ontology (GO) enriched categories for pod transcripts overlapping between WD-, HS-, and WD+HS-altered transcripts (226 transcripts, conserved transcripts from Figure 3c). Abbreviations: BP, biological process; CC, cellular component; HS, heat stress; MF, molecular function; WD, water deficit; WD + HS, a combination of WD and HS.

**Figure S8.**
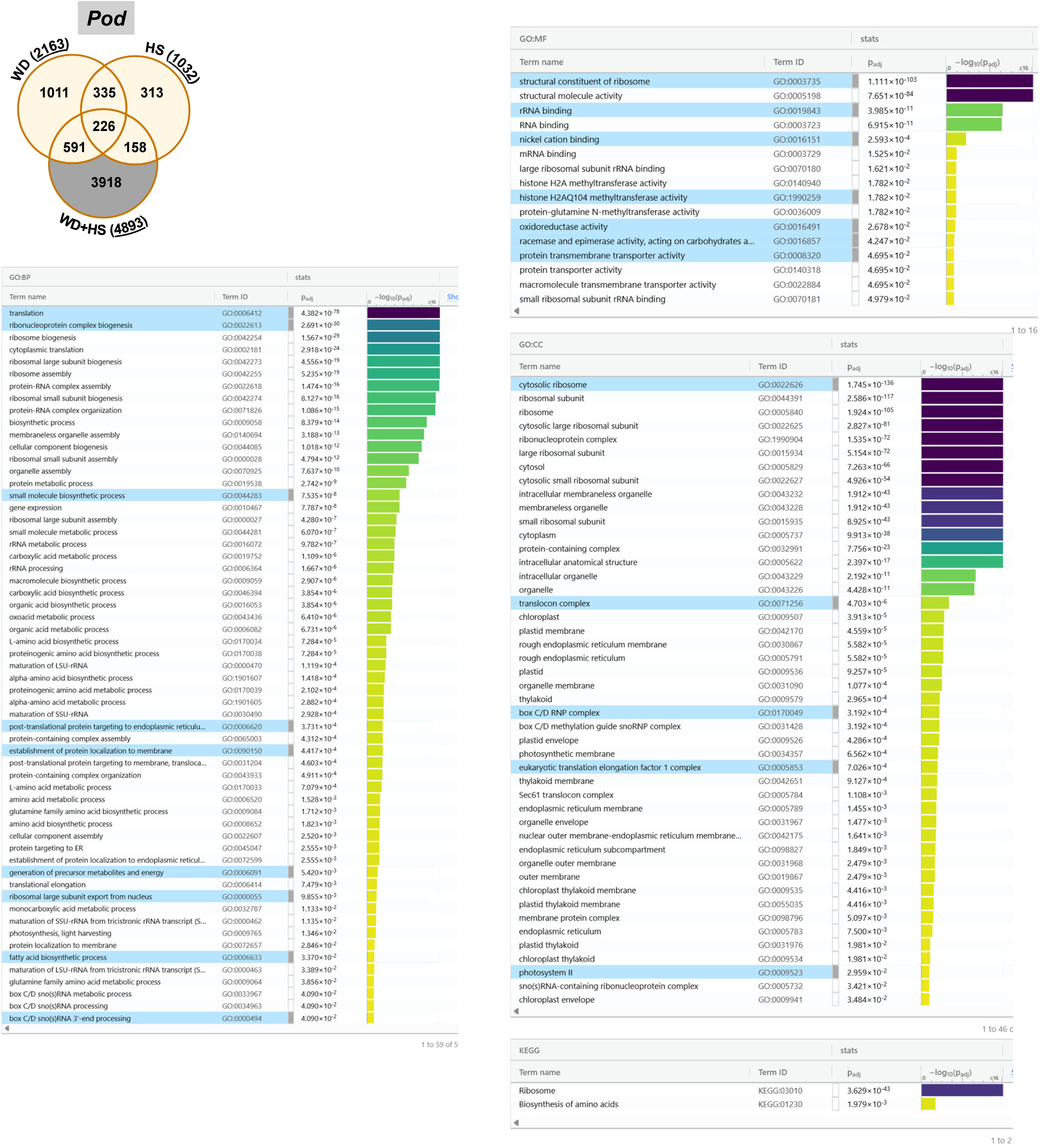
GO enrichment of differentially expressed pod transcripts unique to WD+HS. Gene ontology (GO) enriched categories for pod transcripts uniquely altered under WD+HS stress (3918 transcripts, conserved transcripts from Figure 3c). Abbreviations: BP, biological process; CC, cellular component; HS, heat stress; MF, molecular function; WD, water deficit; WD + HS, a combination of WD and HS.

**Figure S9.**
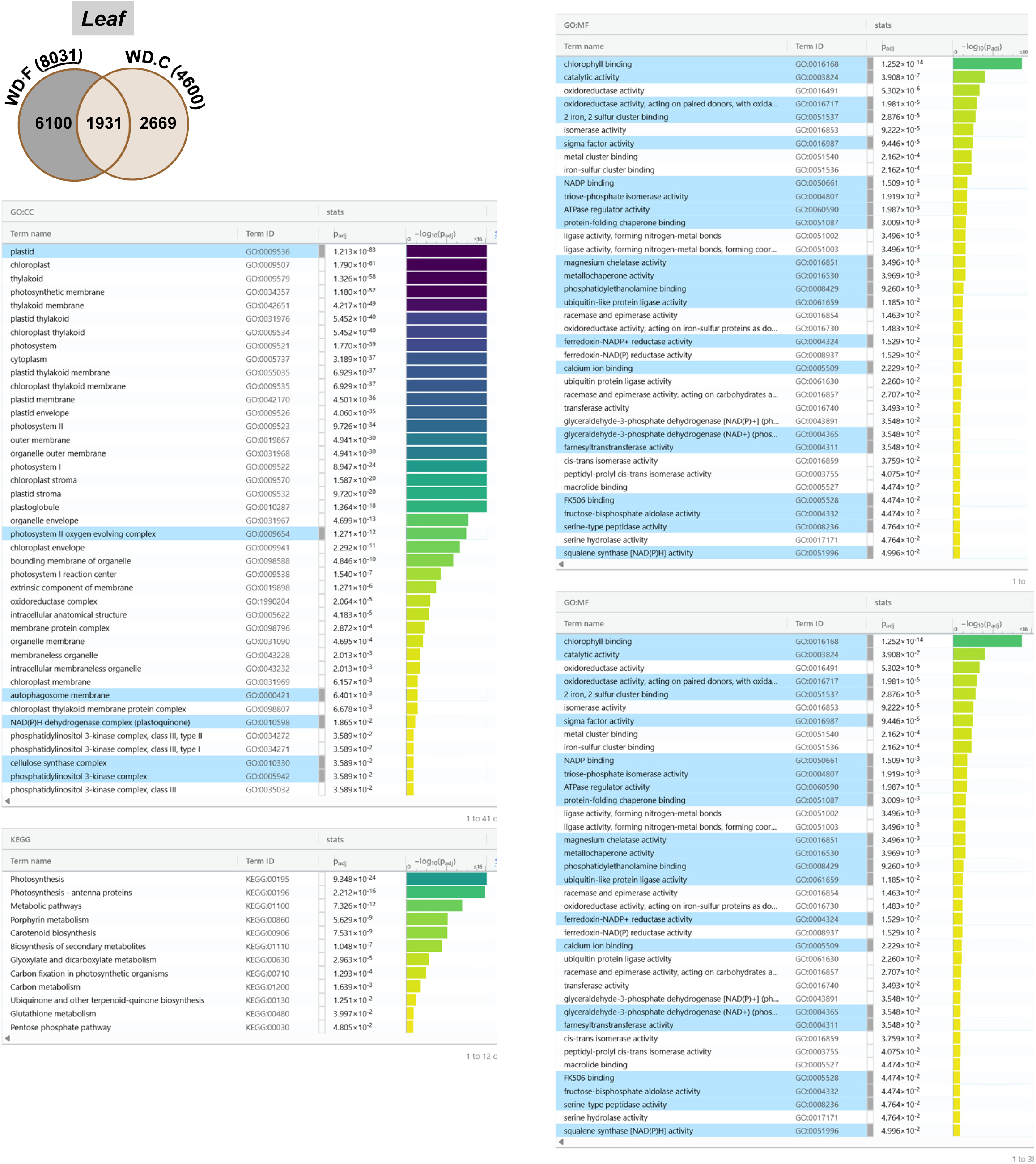
Gene ontology (GO) enriched categories for leaf transcripts uniquely altered in response to WD-stress in field compared to WD-stress in growth chamber. GO enrichment of 6100 transcripts (conserved transcripts from Figure 4a). Abbreviations: BP, biological process; CC, cellular component; MF, molecular function; WD, water deficit.

**Figure S10.**
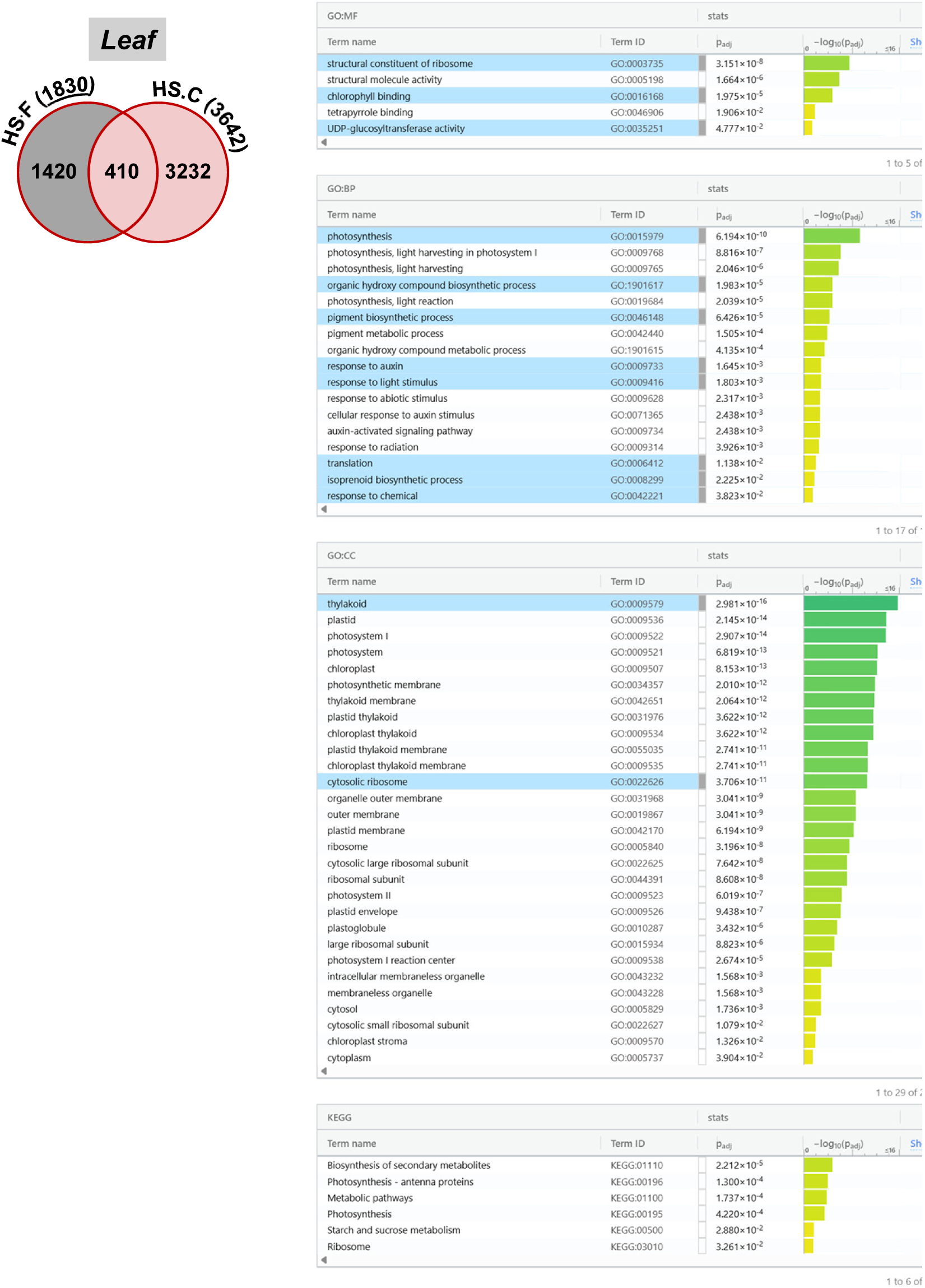
Gene ontology (GO) enriched categories for leaf transcripts uniquely altered in response to HS-stress in field compared to HS-stress in growth chamber. GO enrichment of 1420 transcripts (conserved transcripts from Figure 4a). Abbreviations: BP, biological process; CC, cellular component; HS, heat stress; MF, molecular function.

**Figure S11.**
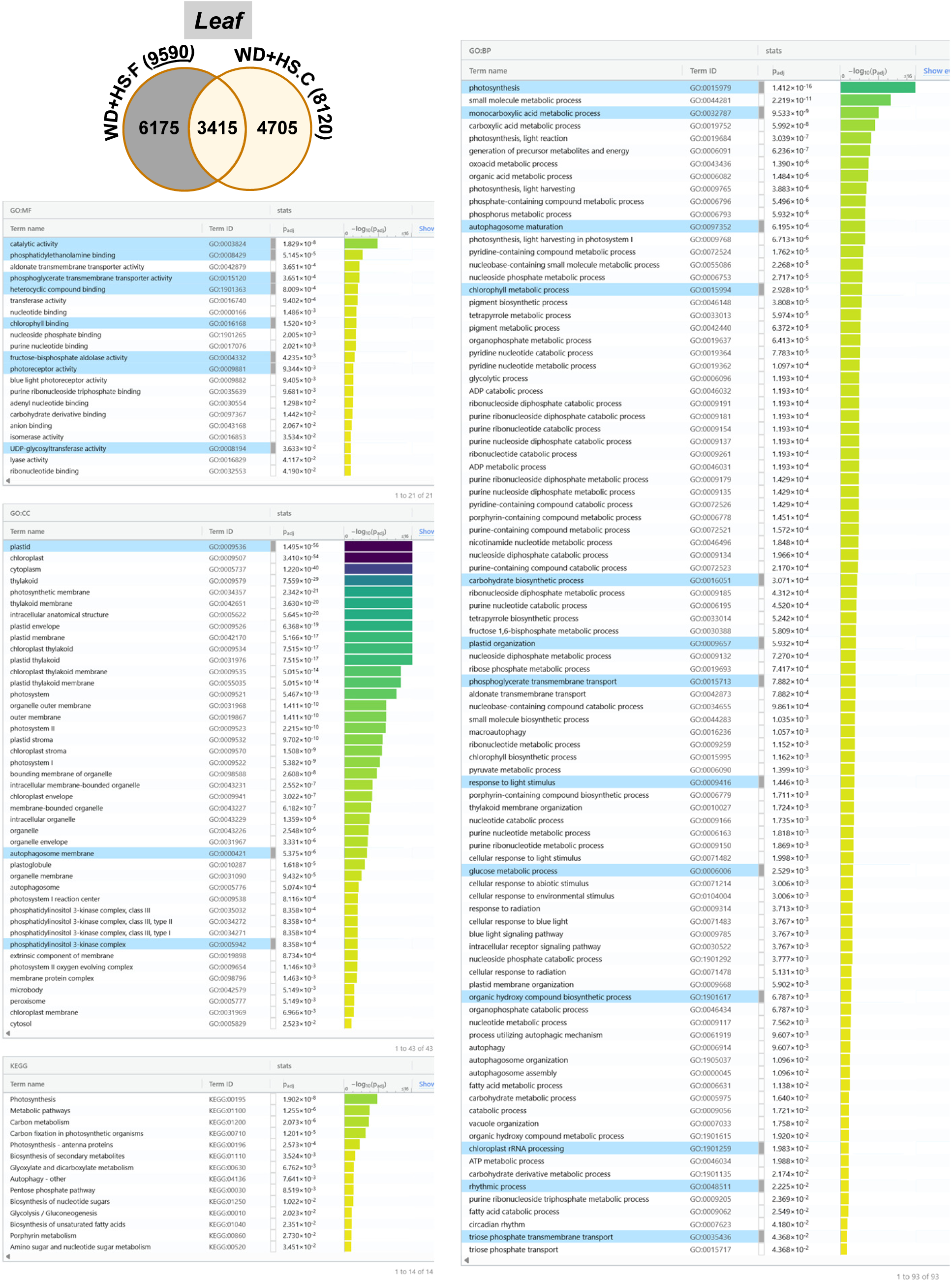
Gene ontology (GO) enriched categories for leaf transcripts uniquely altered in response to WD+HS-stress in field compared to WD+HS-stress in growth chamber. GO enrichment of 6175 transcripts (conserved transcripts from Figure 4a). Abbreviations: BP, biological process; CC, cellular component; MF, molecular function; WD + HS, a combination of WD and HS.

**Figure S12.**
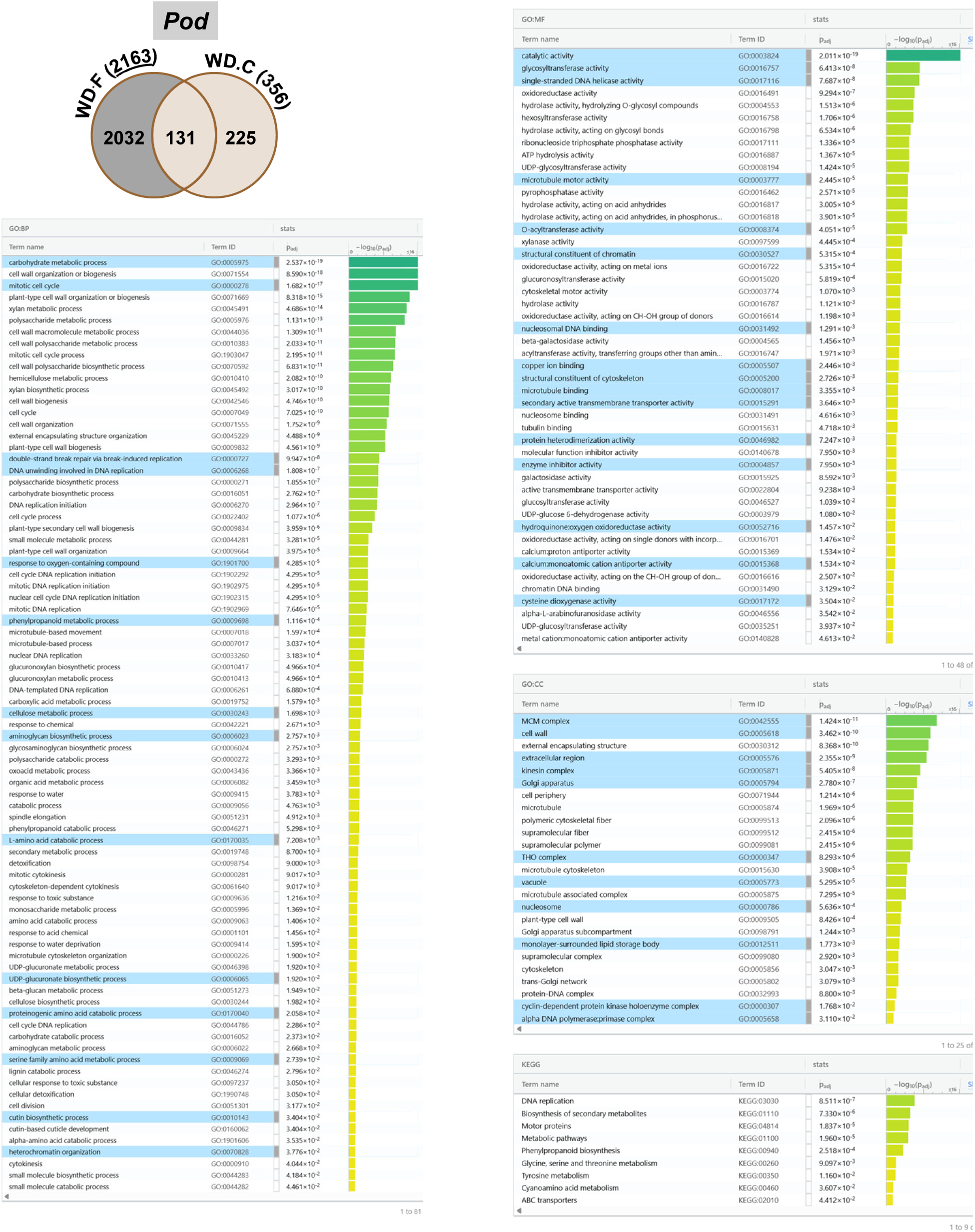
Gene ontology (GO) enriched categories for pod transcripts uniquely altered in response to WD-stress in field compared to WD-stress in growth chamber. GO enrichment of 2032 transcripts (conserved transcripts from Figure 4a). Abbreviations: BP, biological process; CC, cellular component; MF, molecular function; WD, water deficit.

**Figure S13.**
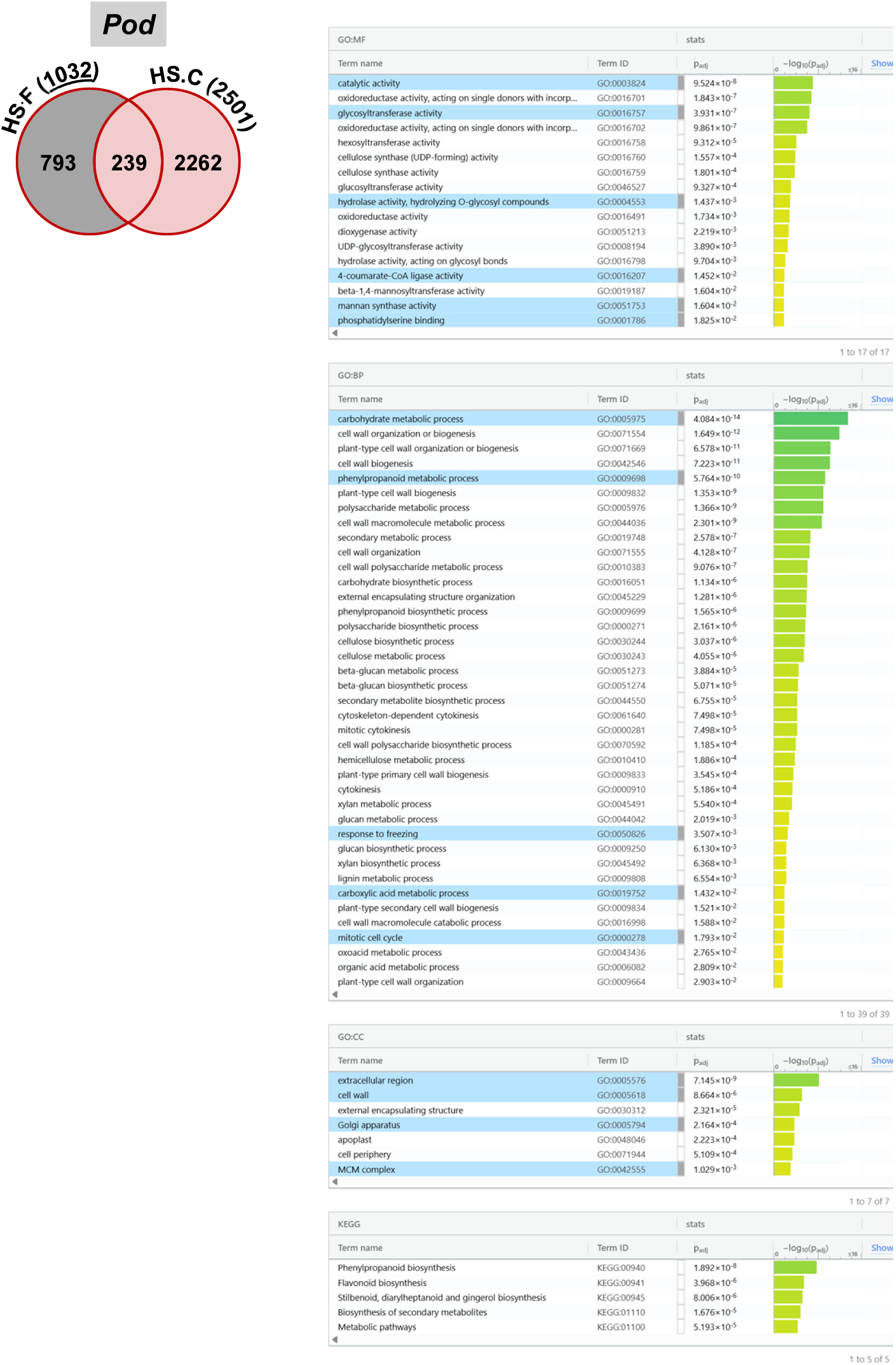
Gene ontology (GO) enriched categories for pod transcripts uniquely altered in response to HS-stress in field compared to HS-stress in growth chamber. GO enrichment of 793 transcripts (conserved transcripts from Figure 4a). Abbreviations: BP, biological process; CC, cellular component; HS, heat stress; MF, molecular function.

**Figure S14.**
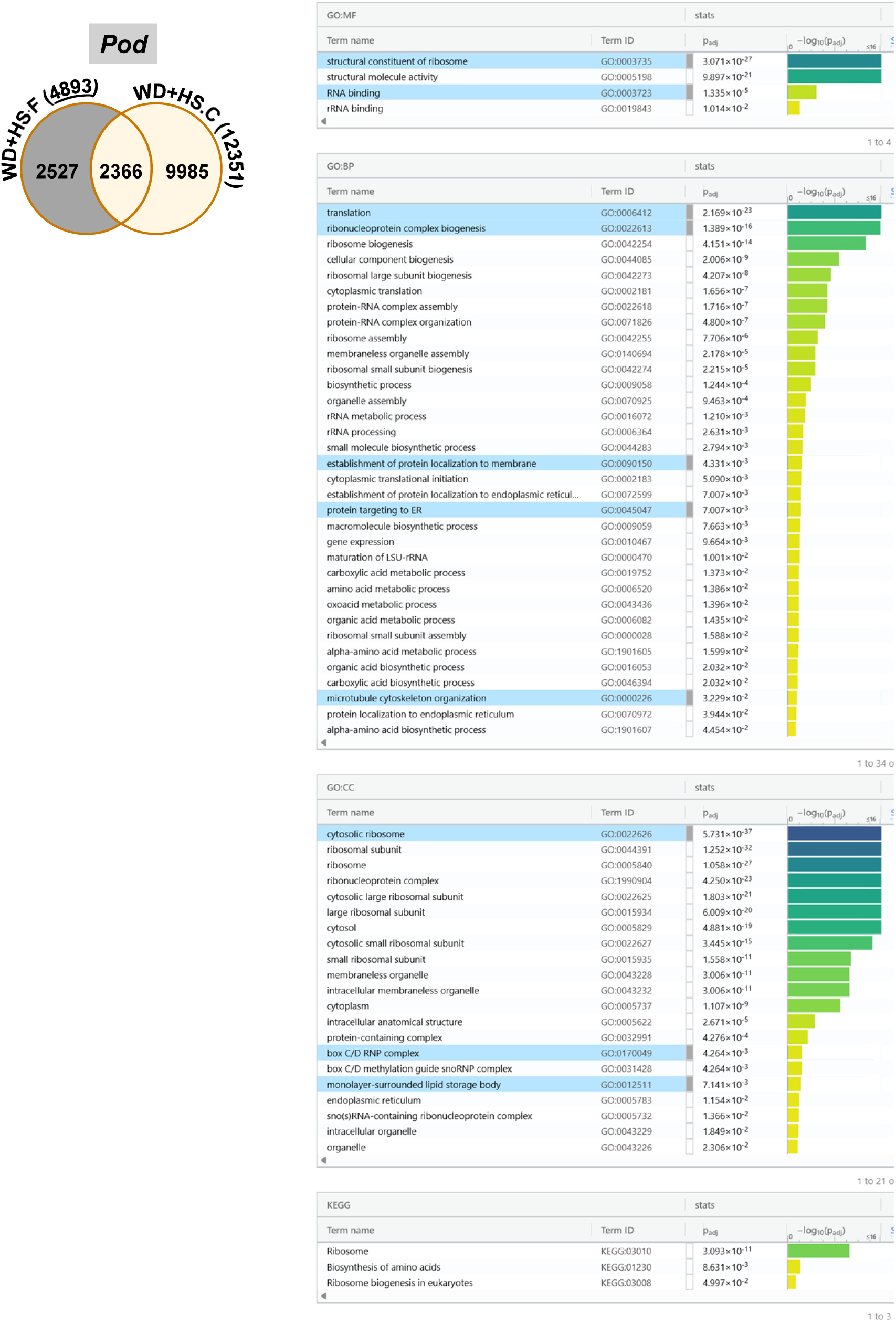
Gene ontology (GO) enriched categories for pod transcripts uniquely altered in response to WD+HS-stress in field compared to WD+HS-stress in growth chamber. GO enrichment of 2527 transcripts (conserved transcripts from Figure 4a). Abbreviations: BP, biological process; CC, cellular component; MF, molecular function; WD + HS, a combination of WD and HS.

**Figure S15.**
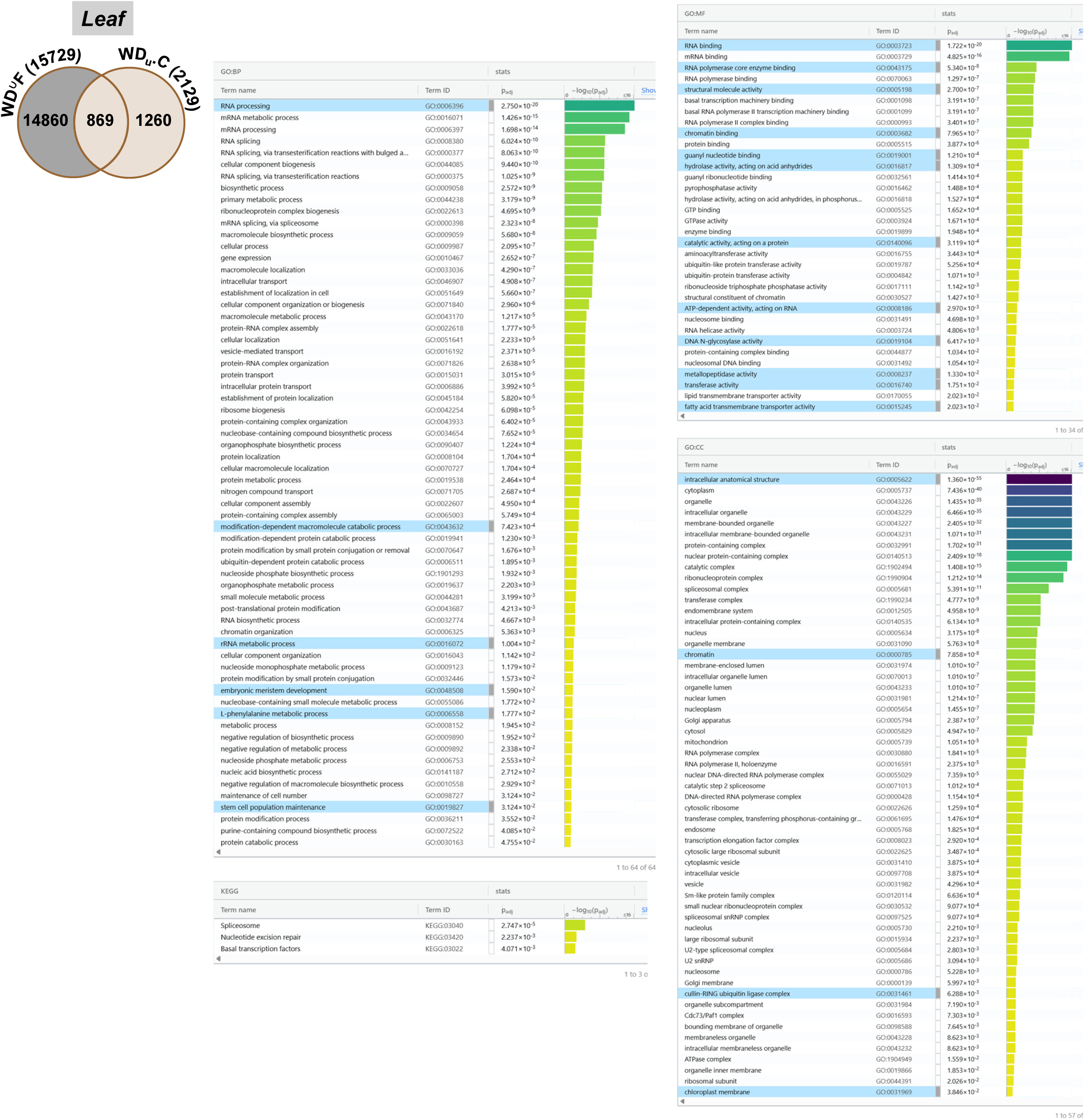
Gene ontology (GO) enrichment of WD induced leaf transcripts unique to field compared to growth chamber. GO enriched categories for leaf transcripts uniquely altered in response to WD-stress in field (unique to each batch year/time pooled together; represented as ^∪^) compared to leaf transcripts unique to WD-stress in growth chamber (14860 transcripts, from Figure 4c). Abbreviations: BP, biological process; CC, cellular component; MF, molecular function; “^∪^”, union of unique genes; “u”, unique to stress treatment; WD, water deficit.

**Figure S16.**
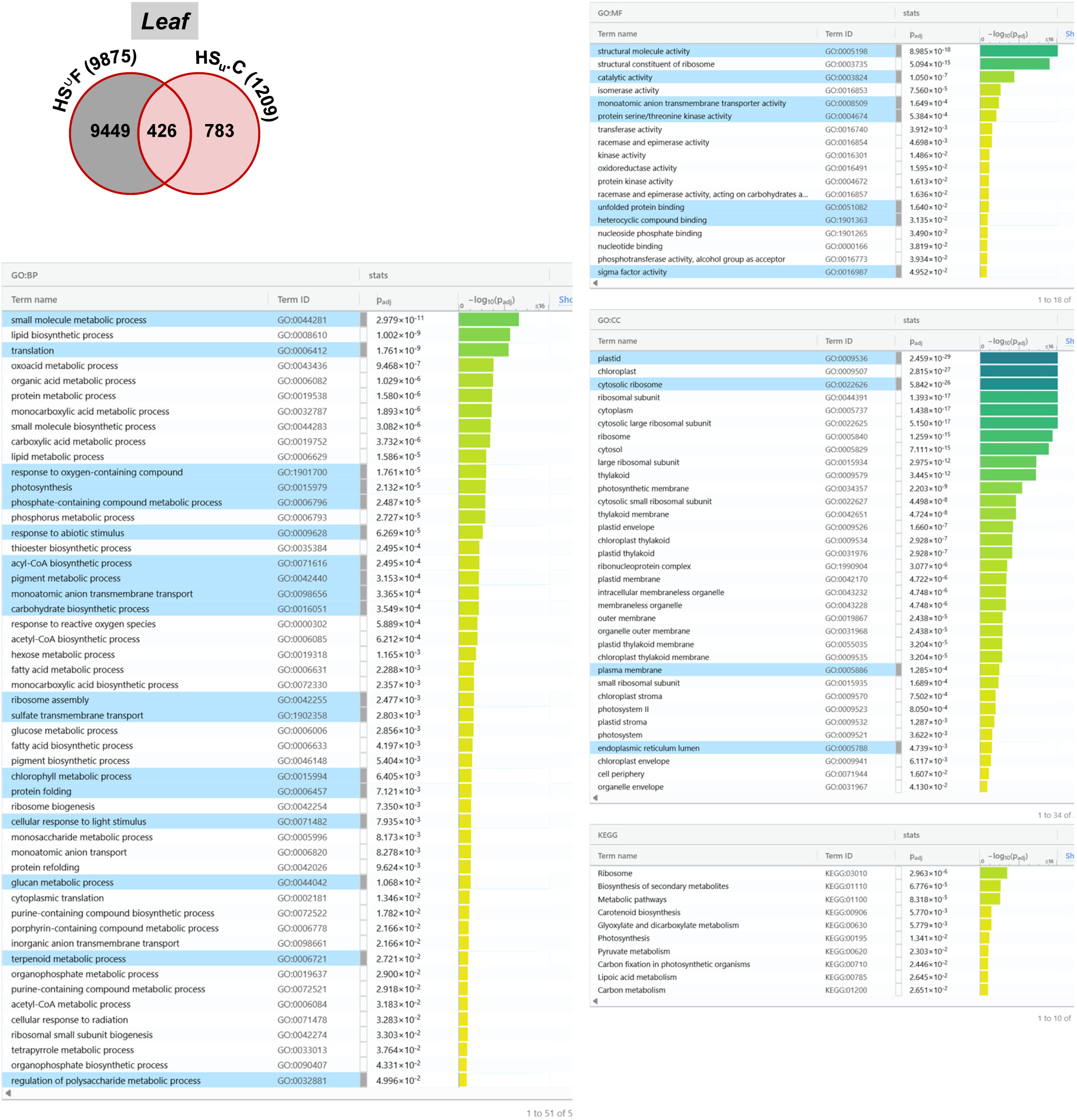
Gene ontology (GO) enrichment of HS induced leaf transcripts unique to field compared to growth chamber. Gene ontology (GO) enriched categories for leaf transcripts uniquely altered in response to HS-stress in field (unique to each batch year/time pooled together; represented as ^∪^) compared to leaf transcripts unique to HS-stress in growth chamber (9449 transcripts, from Figure 4c). Abbreviations: BP, biological process; CC, cellular component; HS, heat stress; MF, molecular function; “^∪^”, union of unique genes; “u”, unique to stress treatment.

**Figure S17.**
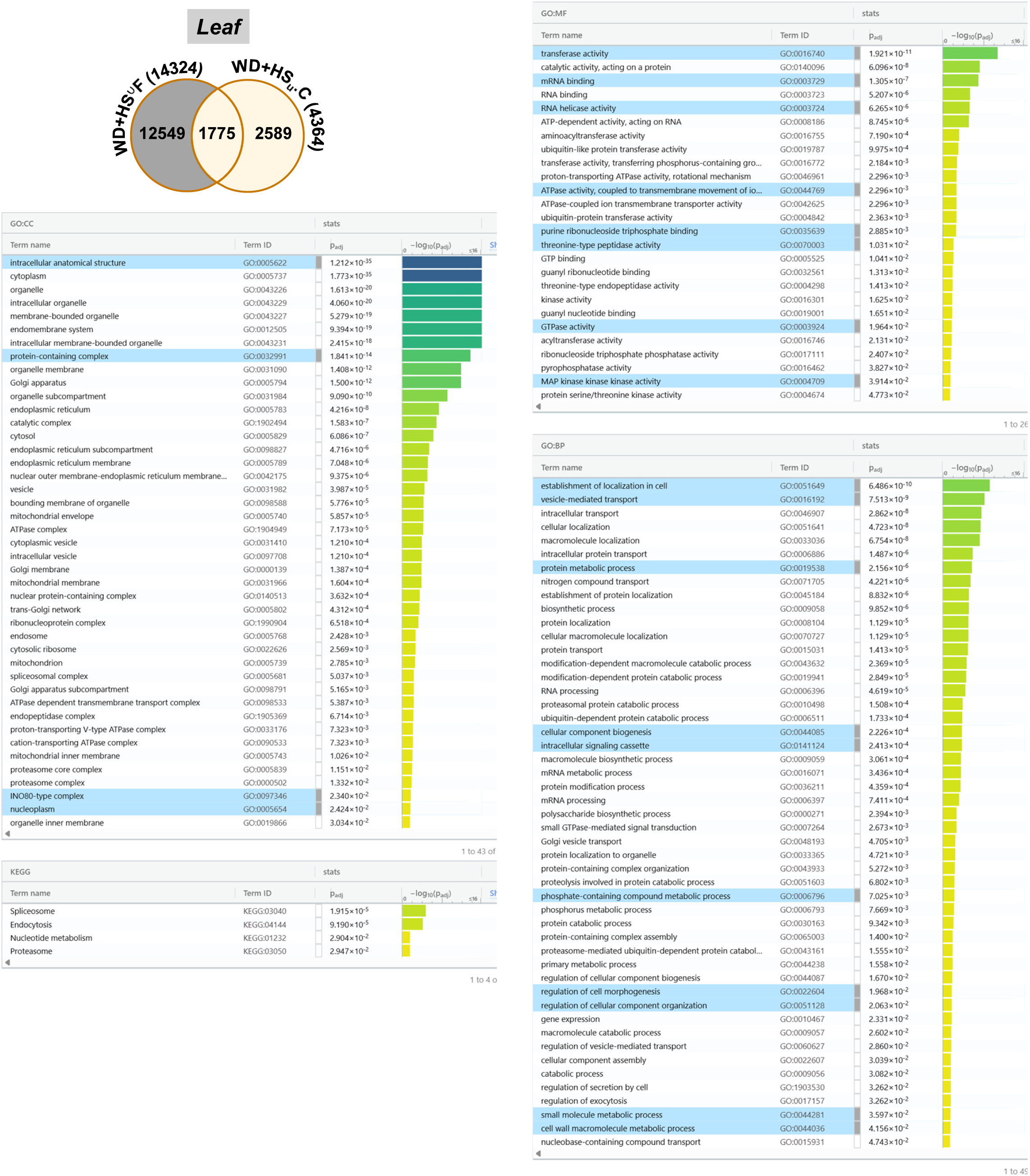
Gene ontology (GO) enrichment of WD+HS induced leaf transcripts unique to field compared to growth chamber. Gene ontology (GO) enriched categories for leaf transcripts uniquely altered in response to WD+HS-stress in field (unique to each batch year/time pooled together; represented as ^∪^) compared to leaf transcripts unique to WD+HS-stress in growth chamber (12549 transcripts, from Figure 4c). Abbreviations: BP, biological process; CC, cellular component; MF, molecular function; “^∪^”, union of unique genes; “u”, unique to stress treatment; WD + HS, a combination of WD and HS.

**Figure S18.**
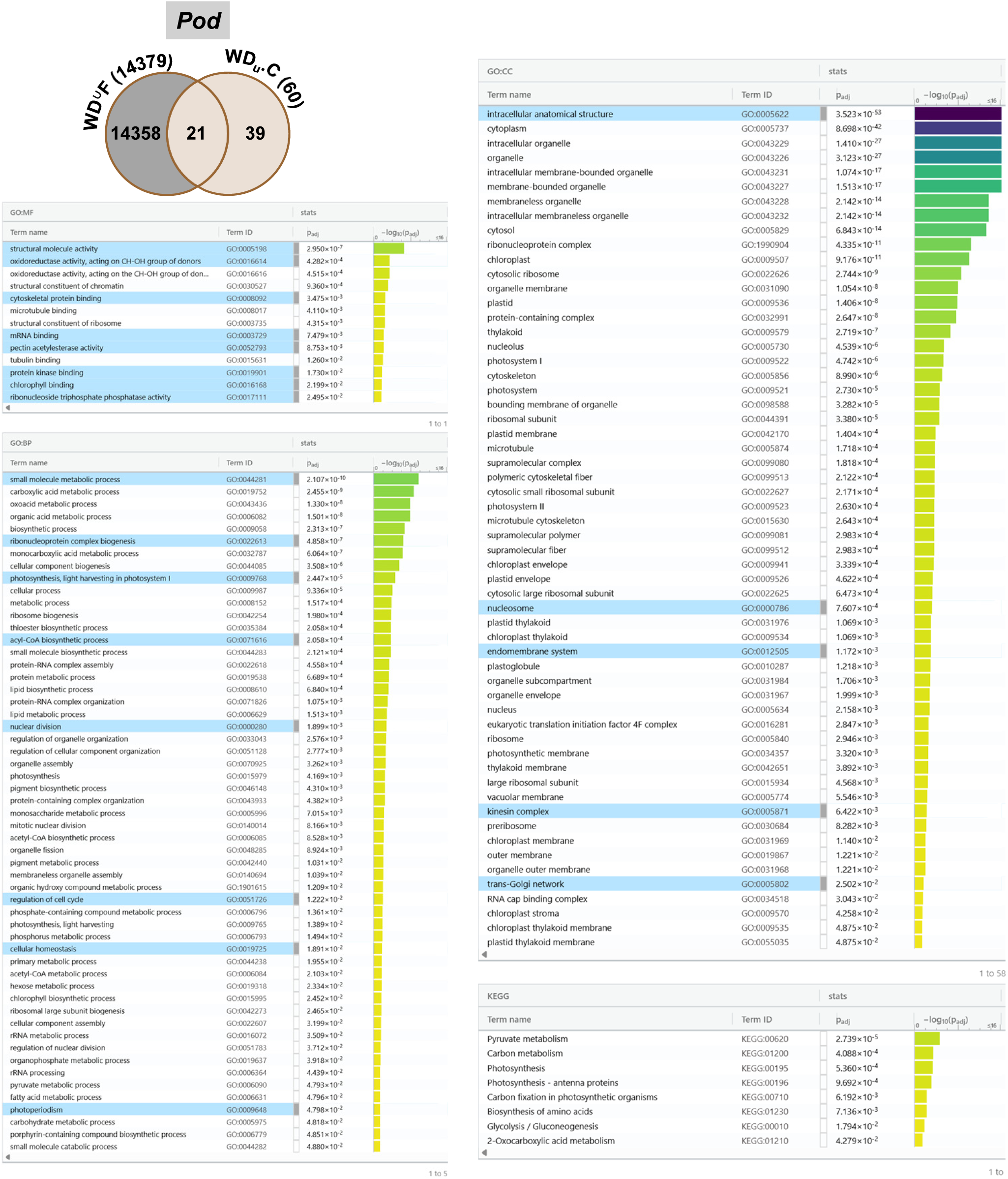
Gene ontology (GO) enrichment of WD induced pod transcripts unique to field compared to growth chamber. Gene ontology (GO) enriched categories for pod transcripts uniquely altered in response to WD –stress in field (unique to each batch year/time pooled together; represented as ^∪^) compared to pod transcripts unique to WD –stress in growth chamber (14358 transcripts, from Figure 4c). Abbreviations: BP, biological process; CC, cellular component; MF, molecular function; “^∪^”, union of unique genes; “u”, unique to stress treatment; WD, water deficit.

**Figure S19.**
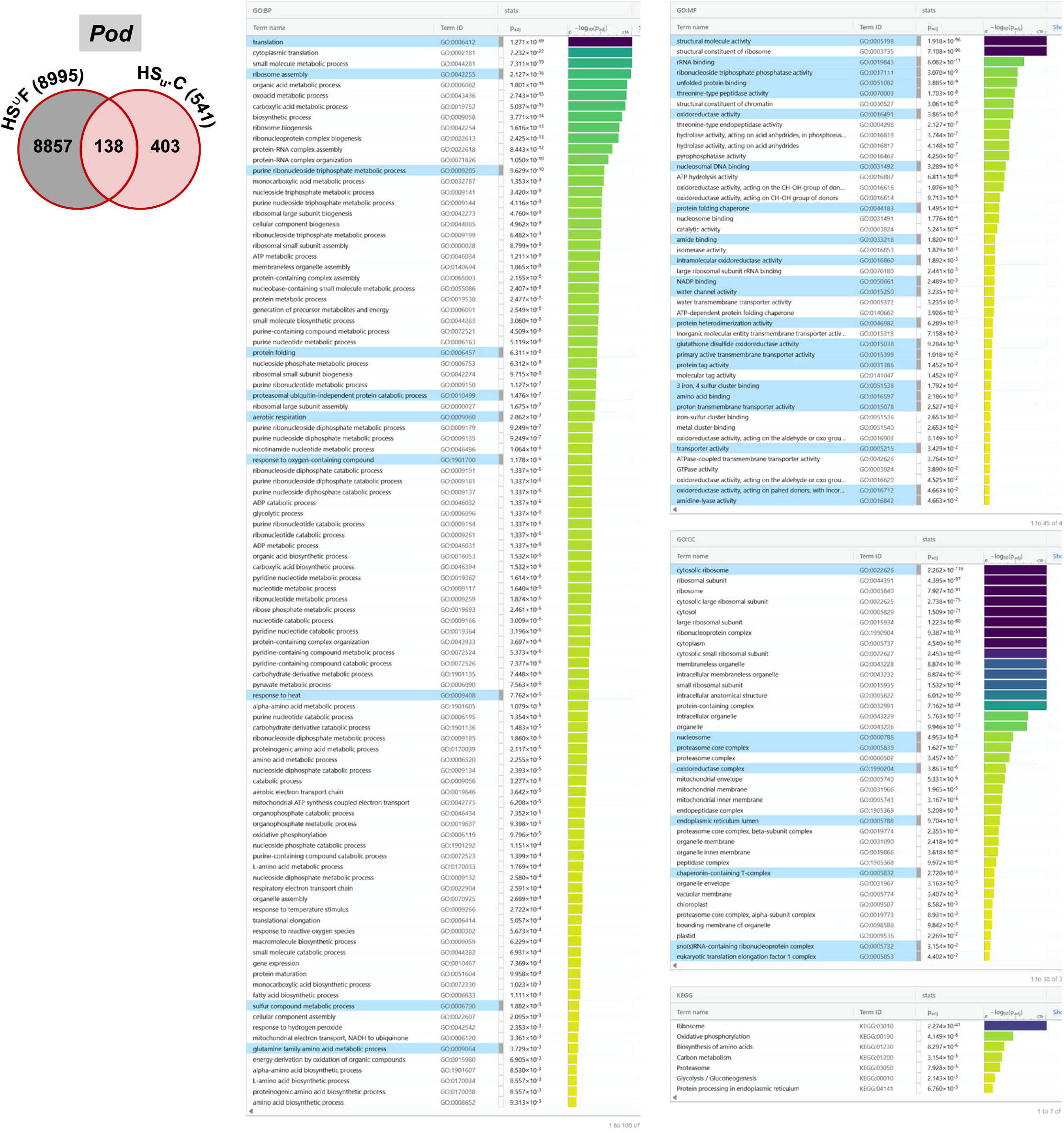
Gene ontology (GO) enrichment of HS induced pod transcripts unique to field compared to growth chamber. Gene ontology (GO) enriched categories for pod transcripts uniquely altered in response to HS-stress in field (unique to each batch year/time pooled together; represented as ^∪^) compared to pod transcripts unique to HS-stress in growth chamber (8857 transcripts, from Figure 4c). Abbreviations: BP, biological process; CC, cellular component; HS, heat stress; MF, molecular function; “^∪^”, union of unique genes; “u”, unique to stress treatment.

**Figure S20.**
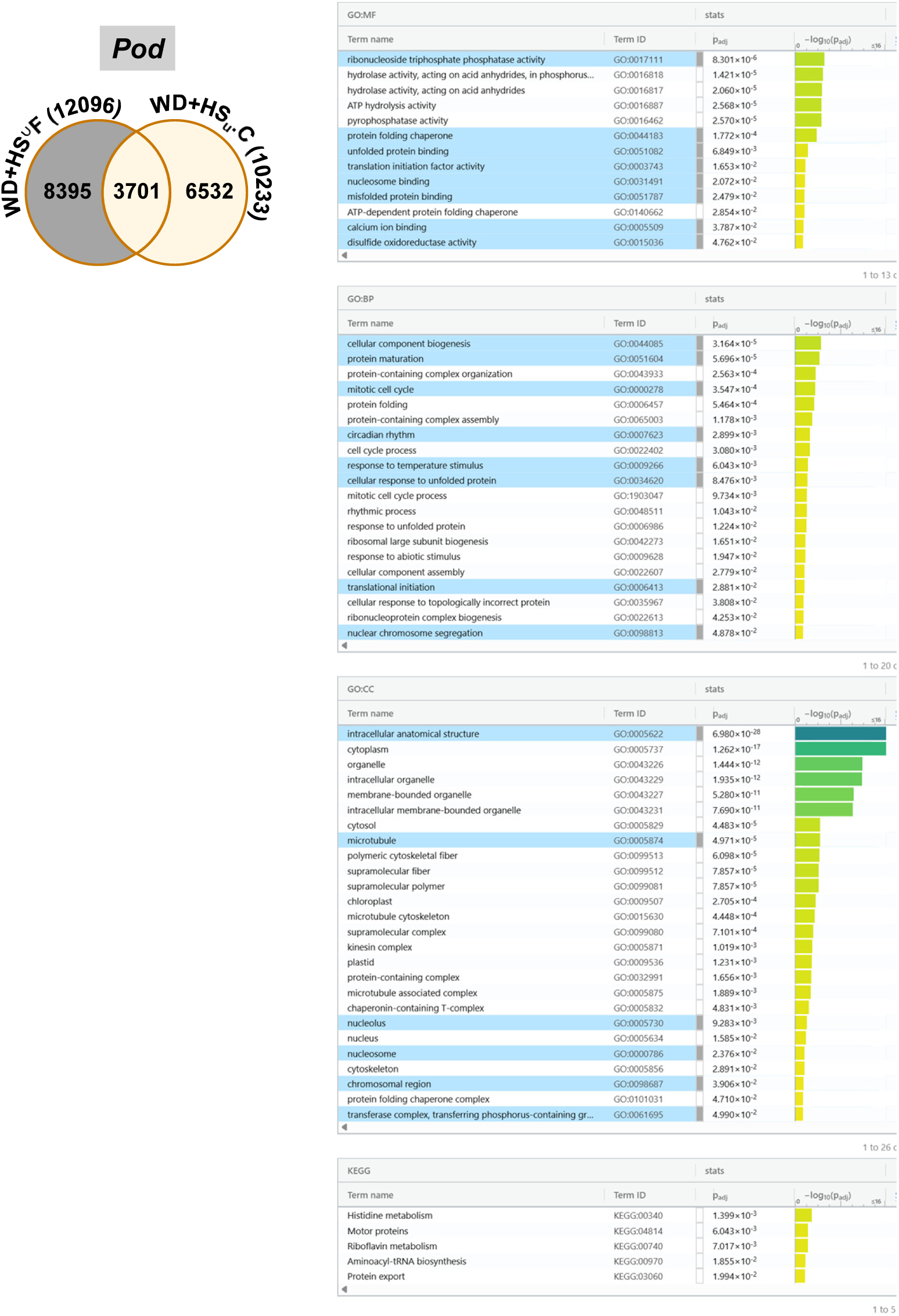
Gene ontology (GO) enrichment of WD+HS induced pod transcripts unique to field compared to growth chamber. Gene ontology (GO) enriched categories for leaf transcripts uniquely altered in response to WD+HS-stress in field (unique to each batch year/time pooled together; represented as ^∪^) compared to leaf transcripts unique to WD+HS-stress in growth chamber (8395 transcripts, from Figure 4c). Abbreviations: BP, biological process; CC, cellular component; MF, molecular function; “^∪^”, union of unique genes; “u”, unique to stress treatment; WD + HS, a combination of WD and HS.

**Figure S21.**
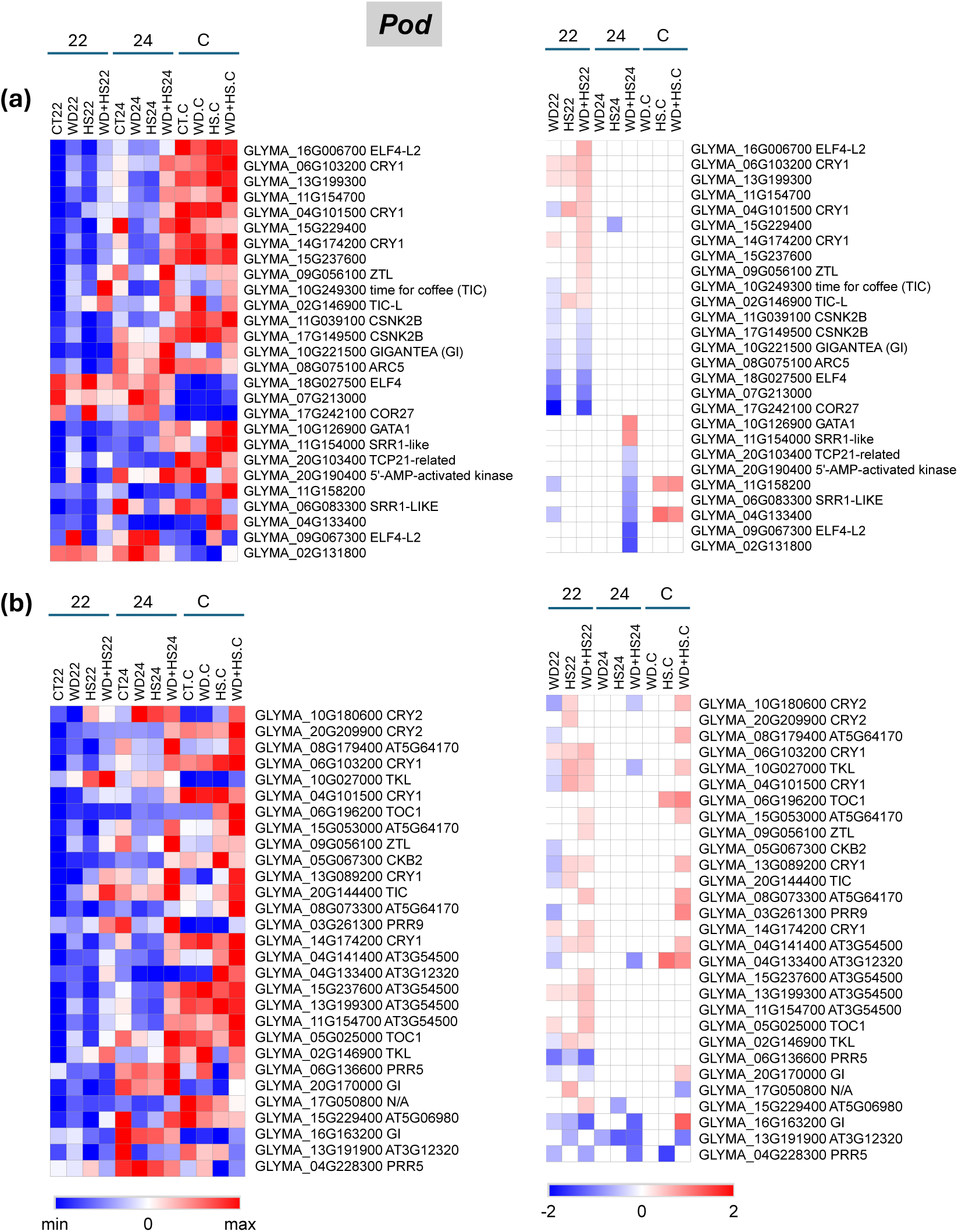
Differential expression of soybean pod transcripts enriched in circadian rhythm and rhythmic processes GO categories. Heat maps showing the differential expression of transcripts encoding factors involved in circadian rhythm (a) and transcripts encoding factors involve in rhythmic process (b), in field and growth chamber grown plants subjected to water deficit, heat stress, and their combination are shown. Significant changes in transcript expression compared to control were defined as adjusted P < 0.05 (negative binomial Wald test followed by Benjamini–Hochberg correction). Abbreviations: 22, 2022 sample; 24, 2024 sample; “.C”, chamber; CT, control; HS, heat stress; WD, water-deficit; WD + HS, a combination of WD and HS.

**Figure S22.**
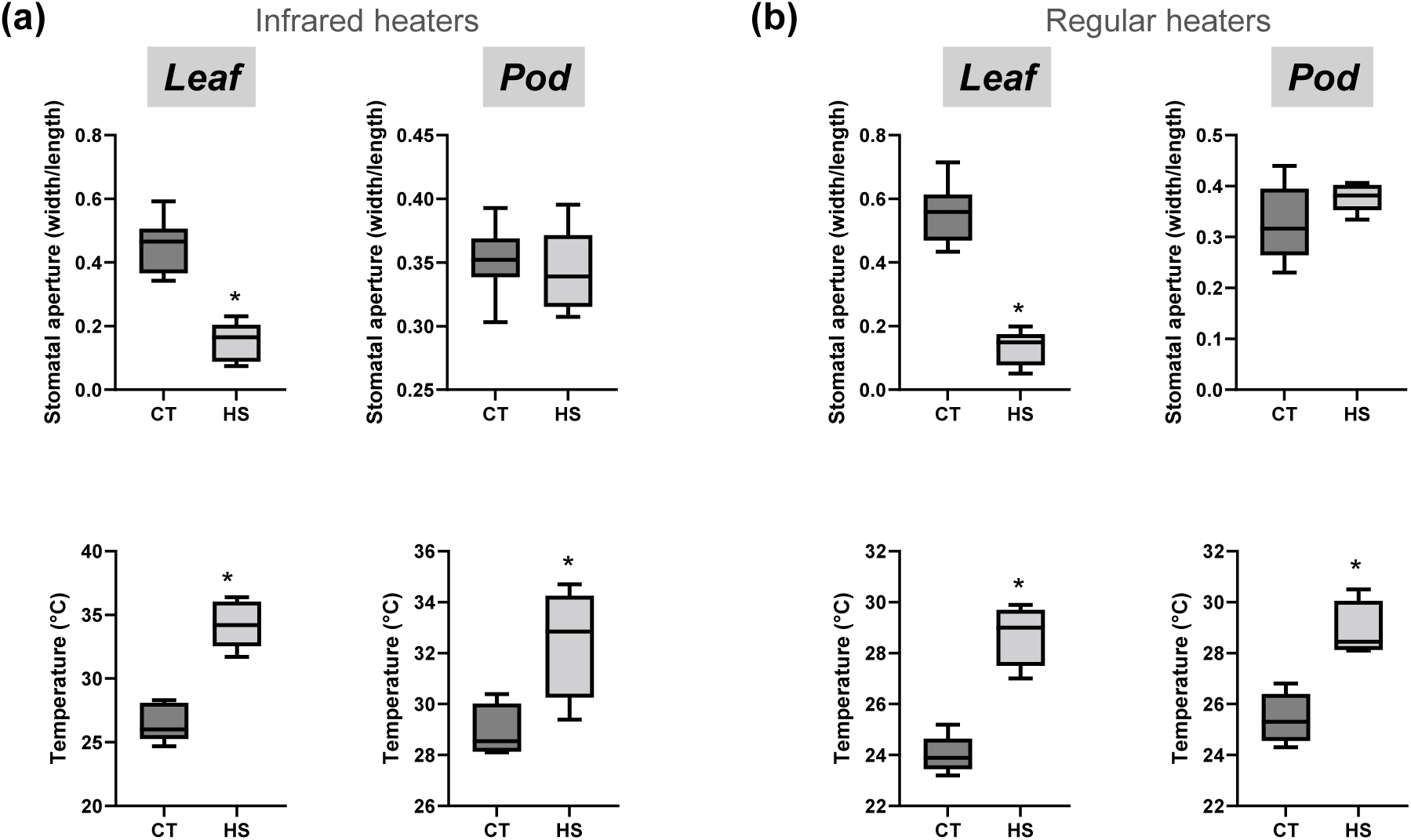
Effects of regular or infrared heaters on stomatal aperture and tissue temperature of pods and leaves. Stomatal aperture of soybean plants grown under control conditions or subjected to heat stress using infrared (a), or space (b) heaters in the field. Stomatal aperture was measured as ratio of width and length for twenty microscopic fields from the middle section of leaves (between the main veins) and from the outer surface of pods (above developing seeds). Box plot represents average of at least 3-5 replicates. Asterisks denote significance at P < 0.05 (Student’s t-test). Abbreviations: CT, control; HS, heat stress.

**Figure S23.**
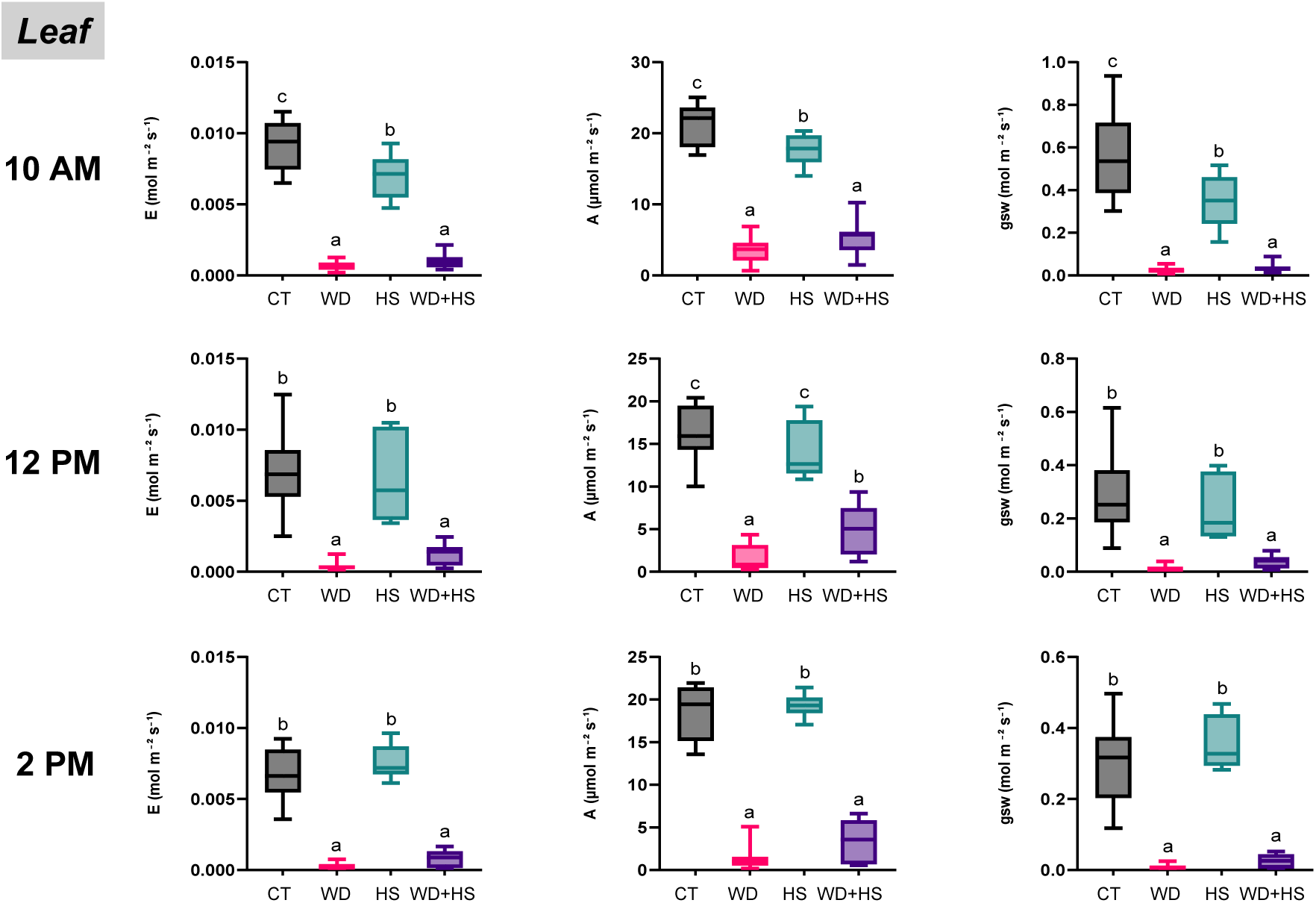
Measurement of gas exchange parameters at different time-of-day in the field. Measurements of leaf photosynthetic activity (A), transpiration (E), and stomatal conductance (gsw) were conducted on soybean plants subjected to water deficit, heat stress, and their combination, at three different times of day. Box plot represents average of at least 3-5 replicates. One-way-ANOVA with Fisher’s least square difference (LSD) was used for test of significance. Different letters denote significant difference at P < 0.05. Abbreviations: CT, control; HS, heat stress; WD, water deficit; WD + HS, a combination of WD and HS.

**Table S1.** Experimental details and environmental parameters.

|  |  |  |  |  |  |  |  |  |
| --- | --- | --- | --- | --- | --- | --- | --- | --- |
|  | **2022** |  |  |  | **2024** |  |  |  |
| Date of seed sowing | May 13<sup>th</sup> |  |  |  | May 23<sup>rd</sup> |  |  |  |
| Initiation of drought stress | June 10<sup>th</sup> |  |  |  | Aug 1<sup>st</sup> |  |  |  |
| Initiation of heat stress | July 25<sup>th</sup> |  |  |  | Aug 19<sup>th</sup> |  |  |  |
| Date collected | July 30<sup>th</sup> |  |  |  | Aug 23<sup>rd</sup> |  |  |  |
| **Environmental condition at the time of sampling** |  |  |  |  |  |  |  |  |
| Air temperature | 24.9 °C |  |  |  | 24 °C |  |  |  |
| Air relative humidity | 59 % |  |  |  | 63 % |  |  |  |
| Average solar radiation (watt/m2) | 650 |  |  |  | 712 |  |  |  |
| Total radiation energy density (KJ/M<sup>2</sup>) | 2339.13 |  |  |  | 2564.36 |  |  |  |
| **Average of environmental condition at the day of sampling** |  |  |  |  |  |  |  |  |
| Maximum air temperature | 26.2 °C |  |  |  | 27.8 °C |  |  |  |
| Average air temperature | 21.50 ± 3.29 °C |  |  |  | 19.77 ± 5.76 °C |  |  |  |
| Average daily air humidity | 63 ± 8.95 % |  |  |  | 76.41 ± 15.29 % |  |  |  |
| Daily average vapor pressure deficit (VPD KPA) | 0.674 |  |  |  | 0.681 |  |  |  |
| Soil water potential (KPA) | CT | -10.93 | ± | 0.15 | CT | -10.09 | ± | 0.13 |
|  | WD | -349.85 | ± | 5.99 | WD | -366.83 | ± | 22.20 |
|  | HS | -96.91 | ± | 9.60 | HS | -45.67 | ± | 1.71 |
|  | WD+HS | -431.24 | ± | 21.90 | WD+HS | -530.23 | ± | 39.34 |
| Soil temperature | CT | 23.76 | ± | 0.65 | CT | 20.82 | ± | 0.52 |
|  | WD | 28.27 | ± | 0.61 | WD | 21.79 | ± | 0.47 |
|  | HS | 23.99 | ± | 0.58 | HS | 21.72 | ± | 0.37 |
|  | WD+HS | 26.93 | ± | 0.92 | WD+HS | 22.40 | ± | 0.49 |

